# High throughput screen in a co-culture model to uncover therapeutic strategies to potentiate the cancer-inhibiting properties of the tumor-stroma in pancreatic cancer

**DOI:** 10.1101/2022.03.11.483991

**Authors:** James Mason, Joshua Cumming, Anna U. Eriksson, Carina Binder, Mitesh Dongre, Cedric Patthey, Margarita Espona-Fiedler, Erik Chorell, Daniel Öhlund

## Abstract

<u>P</u>ancreatic stellate <u>c</u>ells (PSCs) differentiate into multiple subtypes of <u>c</u>ancer <u>a</u>ssociated fibroblasts (CAFs) that modulate disease progression in <u>p</u>ancreatic <u>d</u>uctal <u>a</u>deno<u>c</u>arcinoma (PDAC). CAF subtypes demonstrate functional heterogeneity in tumor development, with conflicting consequences for disease progression. Here, we show that <u>my</u>ofibroblastic CAFs (myCAFs), but not inflammatory CAFs (iCAFs), can act to restrain tumor cell growth in an *in vitro* PDAC model of tumor organoids co-cultured with PSCs. Inhibiting myCAF formation by TGF-β pathway inhibition improved tumor organoid growth, indicating that manipulating the balance of CAF subtypes may be exploited as a therapeutic approach. We therefore conducted a high throughput screen of approximately 36,000 compounds on the co-culture model to find novel compounds to inhibit PDAC tumor cell growth via CAF manipulation. We identify a new role for GNF-5, a known <u>Ab</u>elson tyrosine kinase (Abl) inhibitor, in the context of PDAC as a compound that inhibits tumor cell growth in co-culture; an effect that was accompanied by an induction of the myCAF phenotype around tumor organoids. This highlights the therapeutic potential of novel therapies targeting specific CAF subtypes in PDAC.

## Introduction

Pancreatic cancer, specifically <u>p</u>ancreatic <u>d</u>uctal <u>a</u>deno<u>c</u>arcinoma (PDAC), has one of the poorest survival rates of all cancers with a median survival of approximately 6 months, and a 5-year survival rate that is <11%^1^. Late symptoms, an infiltrative growth pattern, and a lack of accurate and reliable markers for early detection contribute to the fact that metastases are frequently present at the time of diagnosis^2–5^. This prevents surgical treatment, which is the only current/effective curative treatment strategy^4,5^. Unfortunately, for locally advanced and metastatic disease no successful treatment strategy is available, and the tumor soon becomes unresponsive to current chemotherapeutic drugs^6^. PDAC is predicted to become the second leading cause of cancer deaths by 2030^7^ and this is in large part because there is no anticipation for new drugs which can have the potential to drastically improve disease prognosis, and so the need for new treatment strategies is critical.

PDAC is characterized by an abundant tumor stroma accounting for up to 80% of the total tumor mass, containing fibroblasts, immune cells, vasculature, and an <u>e</u>xtra<u>c</u>ellular <u>m</u>atrix (ECM) consisting of predominantly collagens, glycoproteins and glycosaminoglycans^8^. Cancer cells secrete multiple signals into the surrounding tumor <u>m</u>icro<u>e</u>nvironment (TME) that activate nearby fibroblasts, causing them to transdifferentiate into <u>c</u>ancer <u>a</u>ssociated fibroblasts (CAFs)^9–12^. CAFs are important players in PDAC tumor progression by several mechanisms, for instance by providing the cancer cells with soluble and insoluble ligands that stimulate cancer cell growth^13–15^.

A major source of CAFs in PDAC are <u>p</u>ancreatic stellate <u>c</u>ells (PSCs), which are cells normally found in a healthy pancreas^15^. By secreting growth factors such as <u>p</u>latelet <u>d</u>erived <u>g</u>rowth factor (PDGF), fibroblast <u>g</u>rowth factor 2 (FGF-2), transforming <u>g</u>rowth factor <u>b</u>eta (TGF-β) and interleukin <u>1 a</u>lpha (IL-1α) cancer cells stimulate the PSC conversion to CAFs that produce tumor stroma components, generating paracrine loops that promote pancreatic tumor growth^16,17^ (Figure 1a). These paracrine loops present a potential target for novel therapeutics to impede PDAC progression. Indeed, there have been multiple approaches to therapeutically target these loops to treat PDAC but attempts so far have been largely unsuccessful^18–24^. In genetically engineered mouse models of PDAC, general depletion of all CAFs results in poorly differentiated and aggressive tumors which led to worse survival^23,24^, a phenomenon which has also been observed at a clinical trial^22^. These findings together suggest that CAFs have a multifaceted role in PDAC support and progression, and so a more comprehensive knowledge of the role of different subtypes within the tumor microenvironment (TME) is required.

**Figure 1.**
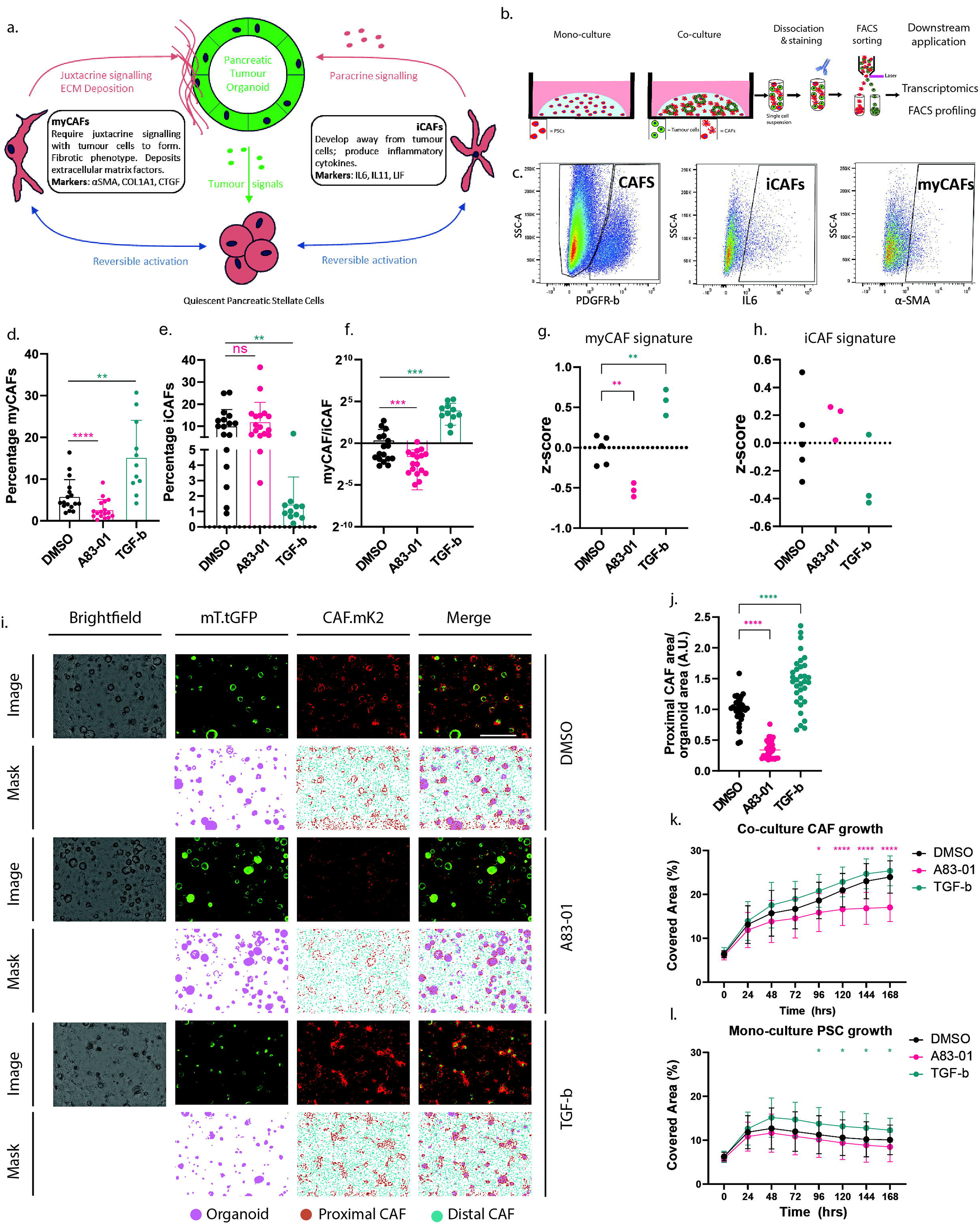
Modulation of CAFs subtype formation and proliferation in a mouse organoid PDAC model. **(A)** Graphical model of cancer cell and <u>p</u>ancreatic stellate <u>c</u>ell (PSC) interactions in <u>p</u>ancreatic <u>d</u>uctal <u>a</u>deno<u>c</u>arcinoma (PDAC). Cancer cells secrete factors that activate resident PSCs in the pancreatic stroma whereupon they form <u>c</u>ancer <u>a</u>ctivated fibroblasts (CAFs), of which there are multiple subtypes. The two best characterized subtypes are <u>my</u>ofibroblastic like CAFs (myCAFs) and inflammatory CAFs (iCAFs). myCAFs are found juxtaposed to cancer cells, requiring juxtacrine signaling for their formation and persistence. myCAFs are readily marked by <u>a</u>lpha smooth <u>m</u>uscle <u>a</u>ctin (α-SMA) and are heavily involved in extra-cellular matrix deposition and remodeling. iCAFs are found distal to the cancer cells and secrete a variety of inflammatory cytokines. The various CAF subtypes are an integral function of tumor progression that influence the cancer cells in this multifaceted, heterogeneous system. **(B)** Illustration of co-culture processing for downstream applications. Here, co-cultures maintained in a three-dimensional matrigel scaffold are enzymatically dissociated into single cell suspensions that are then antibody stained according to the antigen(s) required (for example α-SMA as a marker for myCAFs) followed by flow cytometric analysis and fluorescence-<u>a</u>ctivated <u>c</u>ell sorting (FACS) to isolate cells of interest for further analyses. This approach can be used to profile the myCAF and iCAF complement of a co-culture or may be used to isolate CAF and cancer cell populations from a co-culture for separate transcriptomic profiling. **(C)** Flow cytometry gating strategy for myCAF and iCAF quantification. Co-cultures maintained for five days were treated overnight with a golgi plug then enzymatically dissociated to single cell and stained with antibodies against PDGFR-b, IL6 and α-SMA proteins as well as a live/dead stain and DAPI. Cells were analysed by flow cytometry and gated for size, singlets and being alive (detailed in Supplementary figure 1). Cells were then gated for PDGFR-b expression as a CAF marker, then examined for α-SMA expression (myCAF marker) and IL-6 (iCAF marker). Tumor mono-cultures were used as a negative control for gate determination. **(D-F)** The effect of TGF-β pathway agonists and antagonists on CAF subtype formation and proliferation in mono-culture and in co-culture with PDAC cancer cells was analysed by flow cytometry. Cancer naïve PSCs were co-cultured together with cancer cells and exposed to recombinant TGF-β (2ng/ml, n=11 across three biological replicates), A83-01 (1μM, TGF-β pathway inhibitor, n=14 across three biological replicates) or <u>d</u>i<u>m</u>ethyl sulf<u>o</u>xide (DMSO 1, n=14 across three biological replicates) vehicle control for six days. Cultures were then assessed by flow cytometry for co-culture composition of **(D)** myCAFs, **(E)** iCAFs and **(F)** their myCAF:iCAF ratio. All statistical comparisons performed are Wilcoxon matched pairs signed rank test and conditions are compared against the DMSO vehicle control. **(G)** Bulk transcriptomic signature profiling of 500 CAFs FACS sorted from co-cultures treated for six days with A83-01 (n=3), TGF-β (n=3) or DMSO (n=5) vehicle control. Signatures tested were for transcripts indicative of myCAF and iCAF identity and are plotted as a Z-score. Significance testing performed by one way ANOVA and Dunnett’s multiple comparison test. **(I)** The co-culture model is suited for microscopy-based analyses. Depicted here are fluorescent micrographs of representative co-cultures maintained for 5 days in reduced media conditions in 0.1% DMSO (top), A83-01 (middle) and TGF-β (bottom) and the algorithmic mask detection. Cell lines co-cultured are murine PDAC tumor organoids constitutively expressing turbo-green fluorescent protein (tGFP) and tumor naïve murine PSCs that constitutively express mKate2. Shown in the top panel for each media condition are the brightfield, mT.tGFP (541nm channel), PSC.mKate2 (713nm red channel) and merge of the fluorescent images. The lower panel for each media condition depicts the computationally derived mask of detected objects from each fluorescent channel. From the mT.tGFP channel, detected organoids are highlighted in purple. From the PSC.mKate2 channel, all CAFs are highlighted, and further neural-network trained algorithm identifies “proximal CAFs” (orange) surrounding the detected organoids and the remaining, “distal CAFs” in cyan. All micrographs were taken at the same magnification; scale bar = 1,000μm (top right). **(J-L)** This system was used to analyse **(I)** co-cultures treated with either A83-01 (1μM, n=32 across three biological replicates), recombinant TGF-β (n=33 across three biological replicates) or a DMSO vehicle control (n=32 across three biological replicates) for five days. Calculated is the area of proximal CAFs detected normalized against the area of organoids detected. Significance testing was performed by a two-way ANOVA and a Dunnett’s multiple comparison test, comparing against the DMSO control condition (*, p<0.05; **, p<0.01; ***, p<0.001; ****, p<0.0001). Additionally, these co-cultures were analysed for **(K)** total CAF growth irrespective of subtype. **(L)** Similarly, mono-cultures of PSCs were exposed to TGF-β (n=31 across three biological replicates), A83-01 (n=29 across three biological replicates) or DMSO (n=31 across three biological replicates). Significance testing was performed by a mixed effects model (restricted maximum likelihood method) and a Dunnett’s multiple comparison test, comparing against the DMSO control, where the color of the asterisk refers to the condition being tested. (*, p<0.05; **, p<0.01; ***, p<0.001; ****, p<0.0001).

In addition to promoting tumor growth, stromal elements affect drug response and resistance^25–28^. One example; CD44, a cell surface receptor that is highly upregulated in PDAC, binds hyaluronan which is abundant in PDAC and activates the PI3K-pathway, affecting adhesion, migration, and cancer cell growth^29^. High expression of CD44 correlates to worse prognosis and resistance to Gemcitabine^30^, a standard-of-care drug for PDAC. Enzymatic depletion of hyaluronan reduces tumor stroma, resulting in more efficient drug delivery and a significantly better response to chemotherapy in mice^31,32^, however clinical trials using Pegvorhyaluronidase Alfa (PEGPH20), an enzyme that depletes hyaluronan, have yet failed to improve survival of patients^19^.

Another example is the <u>H</u>edge<u>h</u>og (Hh) signaling pathway, which is active in some CAF subtypes and leads to fibrotic desmoplasia^33^. Inhibition of Hh has proven effective in other cancers such as breast cancer^34^. In the context of PDAC, inhibition of Hh in conjunction with gemcitabine treatment in mice lead to a transient increase in intra-tumoral vascularization, increased intra-tumoral gemcitabine concentration with a consequent transient reduction in tumor growth^35^. Again, these findings were not reflected in a clinical setting, where Hh inhibition provided no added benefit when used in combination with gemcitabine or FOLFIRINOX compared to standard treatment with either drug regimen without Hh pathway inhibition^18,20,21^. Altogether, these studies indicate that the tumor microenvironment, exist in a dynamic state with functional diversity. In turn, this hints that a nuanced approach utilizing targeted depletion of stromal components may be a promising avenue for PDAC therapy.

Activating point mutations in the KRAS proto-oncogene are present in >90% of PDAC patients and point mutations of the tumor suppressor TP53 are described as present in 75% of cases^36–39^. The established “KPC” mouse model (double mutant <u>K</u>ras^+/G12D^ T<u>p</u>53^+/R172H^; expressing Pdx1-<u>C</u>re) harnesses the pancreas-specific Pdx1 promoter to drive conditional mutations in KRAS and TP53. KPC mice spontaneously develop PDAC that mimics the stromal heterogeneity seen in human disease^40^ developing the characteristic pathological distribution of stromal cells which surround the PDAC tumor duct. This is accompanied by high expression levels of <u>a</u>lpha smooth <u>m</u>uscle <u>a</u>ctin (αSMA, encoded by the *ACTA2 gene*). Similarly, we have developed an *in vitro* co-culture model by mixing PSCs and pancreatic ductal organoids derived from primary KPC murine tumors that also recapitulates these aspects of the tumor-stroma^41^. In particular, the co-culture model replicates a symbiotic relationship between the organoids and PSCs, where the PSCs are activated into CAFs and the growth of both tumor and stromal cell types are stimulated^41^.

The co-culture model also generates multiple subtypes of CAFs found in PDAC that are spatially and functionally distinct. Inflammatory CAFs (iCAF) are a subtype that form via paracrine signaling from the tumor cells and are consequently found distal to the tumor cells. iCAFs are characterized by their secretion of inflammatory cytokines such as interleukins 6 and 11^41–43^. Another prominent subtype, ACTA2-expressing <u>my</u>ofibroblastic-like CAFs (myCAFs), are found directly proximal to tumor cells and require juxtacrine signaling from tumor cells coupled with TGF-β pathway activation to form (Figure 1a)^41,44^. It has been previously demonstrated that iCAFs and myCAFs are mutually exclusive in function but that it is possible to revert each back to a quiescent phenotype, indicating that CAF composition in the tumor microenvironment can be modulated^41^.

Given the functional diversity of CAFs in the tumor-stroma, the hypothesis that manipulating the balance of CAF subtypes present in the microenvironment will have therapeutic consequences naturally follows. The *in-vitro* co-culture model is an ideal model for investigating this hypothesis as it generates multiple CAF subtypes with distinct phenotypes that are indicated to have competing effects on tumor cell proliferation^41^. Here, we confirm that the balance between iCAFs and myCAFs can be manipulated in co-culture and investigate resultant effects on tumor cell proliferation. It has been shown that myCAFs show an upregulation of TGF-β response genes and quiescent PSCs can be activated to a myCAF phenotype by the addition of recombinant TGF-β^41^. Therefore, we hypothesized that TGF-β pathway manipulation can be utilized to alter the balance between iCAFs and myCAFs and investigated how that affects tumor cell proliferation.

Extrapolating on the initial hypothesis, we postulated that the co-culture model could be used to identify novel therapeutics that disrupt the symbiosis between the tumor cells and CAFs. The nature of the co-culture model; several cell types that are subject to multiple feedback loops, is inherently heterogeneous. Therefore, we tested whether high content imaging analysis of co-cultures allows for measures that could describe, in a multivariate model, usual co-culture growth in the absence of chemical perturbation. These measures were contrasted against co-cultures that were subject to TGF-β pathway inhibition to test whether changes in co-culture composition could be detected. Based upon the results, we proceeded to screen a library of chemical compounds to identify novel therapeutics that alter the balance of CAF subtypes. Thus, over 36,000 compounds from the <u>C</u>hemical <u>B</u>iology <u>C</u>onsortium <u>S</u>weden’s (CBCS, SciLifeLab, Sweden) screening library were tested on the co-culture model.

Overall, using a high throughput screen which combines high content and multivariate analysis in tumor-organoid co-culture models, we demonstrate that CAF subtypes can be manipulated using targeted approaches, and that a shift in balance between myCAFs and iCAFs leads to inhibited tumor cell proliferation in co-culture. We uncover a novel role for GNF-5, a selective allosteric <u>Ab</u>elson tyrosine kinase (Abl) inhibitor^45^, in the PDAC context by inducing a myCAF phenotype around tumor organoids in co-cultures. This highlights the potential of therapies targeting tumor-CAF interactions as a promising approach to treat PDAC.

## Results

### CAF subtype formation can be modulated in co-culture

We utilized a co-culture model of KPC <u>m</u>urine pancreatic ductal tumor organoids (mT) together with tumor-naïve PSCs to examine the interactions between these cell types (Figure 1a). First, we assessed the malleability of CAF subtype formation in co-culture through transcriptomic profiling and flow cytometric analysis when modulating the TGF-β signaling pathway (Figure 1b). Co-cultures were maintained for six days in Matrigel with a basal media supplemented with either an inhibitor of TGF-β type I receptor (A83-01), recombinant TGF-β, or <u>d</u>i<u>m</u>ethyl sulf<u>o</u>xide (DMSO) as a vehicle control. CAFs from these co-cultures, identified by PDGFR-β expression, were then analysed by flow cytometry for <u>a</u>lpha smooth <u>m</u>uscle <u>a</u>ctin (α-SMA) and interleukin 6 (IL-6) as markers for myCAF and iCAFs, respectively (Figure 1a, b, c, Supplementary figure 1). Recombinant TGF-β significantly induced a myCAF phenotype in co-culture (DMSO mean 5.39% vs. recombinant TGF-β mean 15.28%, p=0.002, Figure 1d). In contrast, A83-01 potently inhibits myCAF formation (DMSO mean 5.39% vs. A83-01 mean 2.76%, p<0.0001, Figure 1d) which together confirms that myCAF identity is supported by TGF-β signaling.

With respect to iCAFs, recombinant TGF-β inhibits iCAF formation in co-culture as measured by Il-6 expression by flow cytometry (DMSO mean 10.28% vs. recombinant TGF-β mean 1.50%, p=0.002, Figure 1e), which altogether significantly increases the myCAF:iCAF ratio of the co-cultures (DMSO mean ratio 1.63 vs. recombinant TGF-β mean ratio 16.12, p=0.001, Figure 1f). A83-01 shows no effect on iCAF formation (DMSO mean 10.25% vs. A83-01 mean 12.19, p=0.087, Figure 1e), however its addition results in a significant overall reduction in myCAF:iCAF ratio in co-cultures (DMSO mean ratio 1.34 vs. A83-01 mean ratio 0.3, p=0.0002, Figure 1f).

We further corroborated the effect of TGF-β pathway manipulation on co-cultures by bulk transcriptomic analysis on tumor cells and PSCs from co-cultures treated with A83-01 and TGF-β. The relative change in these signature scores was examined between cultures treated with A83-01 or TGF-β from a DMSO vehicle control. The transcriptomic analyses confirm the alteration in CAF identity where A83-01 significantly suppresses a myCAF phenotype compared to co-cultures in DMSO (p=0.0047, Figure 1g) with a mild, but non-significant induction of an iCAF signature score (Figure 1h). In contrast, recombinant TGF-β strongly induced a myCAF signature score (p=0.0015, Figure 1g) and a mild reduction in the iCAF signature score (Figure 1h).

We next wanted to examine CAF subtype identity and co-culture growth dynamics over time by fluorescence microscopy using mT and PSC cell lines lentivirally modified to constitutively express turboGFP (mT.tGFP) and mKate2 (PSC.mKate2), respectively (Figure 1i, Supplementary Figure 2a,b). Thus, co-cultures were analysed by high content imaging to detect both mT.tGFP organoids and PSC.mKate2-derived CAFs. Further, it has previously been demonstrated that there is a spatial distinction between CAF subtypes, where myCAFs are juxtaposed to tumor cells, since they require juxtacrine signaling to form, and iCAFs are found distal to the tumor cells, responding to paracrine signaling from the tumor cells^41^. Therefore, we utilized an image-based analysis approach to examine the formation of the CAF subtypes in co-culture based upon their spatial relationship to the organoids. Since myCAFs are found directly proximal to tumor cells, CAFs directly proximal to tumor cells are interpreted as likely myCAFs (“Proximal CAFs”, Figure 1i). On the other hand, CAFs distal to tumor cells in these co-cultures are normally comprised of a prominent representation of iCAFs mixed with other CAF subtypes and thus can likely be identified as non-myCAFs^41^ (“Distal CAFs”, Figure 1i). The classification of CAFs as either “proximal CAFs” or “distal CAFs” was performed by neural network-based machine learning algorithms which allowed object identification and classification trained by manual object selection and rejection according to the manufacturer’s guidelines (SoftmaxPro 7, Molecular Devices, provided in online supplementary materials).

Given the juxtacrine requirement for proximal CAF classification, the opportunity of proximal formation is dependent on the availability of cancer organoids. Therefore, an examination of proximal CAF area per available organoid area was examined in co-cultures treated with A83-01 and TGF-β. Addition of A83-01 very significantly reduced the proximal CAF:organoid area ratio in co-cultures (p<0.0001, Figure 1j). In contrast, supplementation of recombinant TGF-β significantly increases the proximal CAF:organoid area ratio (p<0.0001, Figure 1j). Such findings correlate proximal CAF formation with the myCAF formation as determined by flow cytometry (Figure 1d), indicating that “proximal CAF” image-based detection mirrors myCAF formation.

TGF-β pathway manipulation also affects CAF proliferation as well as subtype identity, where the addition of A83-01 significantly inhibits their growth in co-culture (Figure 1k). However, the inhibitory effect of A83-01 on proliferation is not seen in PSC in mono-culture, but instead their growth is significantly increased in the presence of TGF-β (Figure 1l), an effect that is not significant in co-culture (Figure 1k). Additionally, PSC growth is much reduced in each case than when co-cultured with mT cells suggesting potent PSC proliferation is induced by activation in the presence of tumor cells (Figure 1l). Altogether, these data confirm that CAF subtype identity and proliferation can be significantly modified by pharmacological intervention.

### Manipulating CAF fate alters cancer cell behaviour

Given that there are established interactions between CAFs and cancer cells, we explored the effects of manipulating CAF phenotype in co-culture on cancer organoids. These effects were tested in three different biological replicate co-culture combinations and assessed by fluorescent imaging. In each combination, the effect of A83-01 supplementation in the media significantly increased cancer organoid growth in co-culture from between 72-96 hours onwards (Figure 2a). This effect is not seen in tumor mono-cultures treated with A83-01, where instead A83 reduced cancer organoid growth (Figure 2b), indicating that its effect on increasing tumor growth in co-culture is mediated via the CAFs in culture.

**Figure 2.**
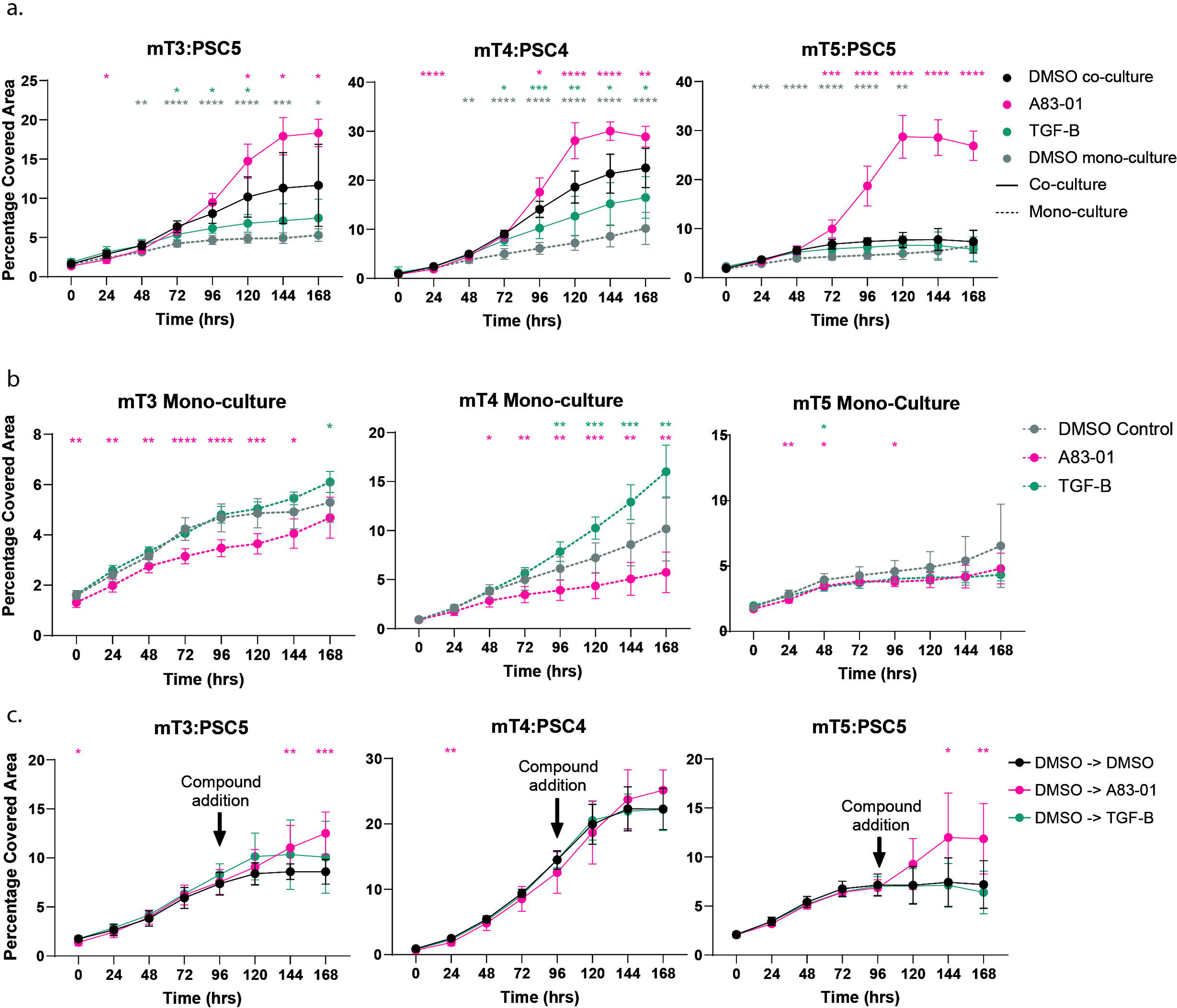
Effect of TGF-β pathway manipulation on cancer cell proliferation. (A, B) Proliferation curves for different biological replicate co-cultures (solid lines) of fluorescent murine pancreatic cancer organoids with PSCs and mono-cultures (dotted line) of fluorescent murine pancreatic cancer organoids determined by image analysis of percentage well coverage of cancer organoids (by fluorescence) over time when exposed to media supplemented with recombinant TGF-β (green, 2ng/ml, n= 33 across three biological replicates), A83-01 (pink, 1uM, n=32 across three biological replicates) or a DMSO (black, 0.1%, n= 33 across three biological replicates) vehicle control. **(C)** Pharmacological intervention on co-cultures where cultures were all treated in media supplemented with DMSO for four days, at which point cultures were supplemented with media containing DMSO (Black, n= 31 across three biological replicates), A83-01 (pink, final concentration 1μM, n= 32 across three biological replicates), or TGF-β (green, final concentration 2ng/ml, n= 32 across three biological replicates). All significance testing was performed by a mixed effects model (restricted maximum likelihood method) and a Dunnett’s multiple comparison test, comparing against the DMSO control, where the color of the asterisk refers to the condition being tested (*, p<0.05; **, p<0.01; ***, p<0.001; ****, p<0.0001).

The converse tends to hold true for the addition of recombinant TGF-β to co-cultures. TGF-β addition can reduce cancer organoid growth in co-culture (Figure 2a), whereas in cancer organoid mono-cultures, their growth is generally increased (Figure 2b). However, the effect of reduced cancer organoid proliferation in co-culture by the addition of recombinant TGF-β appears to be dependent on the cell and organoid lines, since although the mT5:PSC5 combination demonstrated a reduction in cancer cell growth, this effect was not significant. In conclusion, the effect on cancer organoid proliferation is dependent on the interaction between co-cultured CAFs and TGF-β pathway modulation. Given that myCAF formation is regulated through the TGF-β pathway (Figure 1c,d,e), we provide evidence that myCAF prevalence influences cancer organoid proliferation negatively in co-culture.

We next wanted to examine whether the effects of TGF-β pathway augmentation would manifest in co-cultures that were already established rather than with chronic exposure from the time of co-culture seeding. Here, all conditions were initially exposed to a DMSO vehicle control, allowing both iCAFs and myCAFs to form, until media was supplemented at the 96-hour timepoint with the relevant compound (Figure 2c). In each case, the addition of A83-01 induced an increase in cancer cell proliferation within 48 hours, however the supplementation of recombinant TGF-β appeared to have little effect. Altogether, these data demonstrate that cancer cell proliferation can be modulated by manipulation of the surrounding CAFs, an effect that holds true when PSCs have already been established as myCAFs or iCAFs, indicating avenues for targeted tumor-stroma therapies.

### Screening a compound library against a co-culture model

Having identified that cancer cell behaviour can be modulated indirectly by manipulation of the surrounding CAFs, we sought to use co-cultures as a model for use in a high throughput screen to identify compounds that can inhibit cancer cell growth either directly, via, or despite CAF presence. Consequently, we employed a high content image-based analysis capable of effective characterization of the co-cultures including CAF subtype information as described in Figure 1i. An additional analysis was developed with an Arrayscan microscope that possessed greater magnification that could be used in conjunction with the minimax analysis (Supplementary figure 3a). Finally, cultures were analysed by a resazurin reduction assay for NADPH as a readout for their overall metabolic potential. Thus, in these co-cultures, it was possible to examine the co-culture composition in response to small molecule addition.

Therefore, a high throughput screening approach was used to test these co-cultures against a compound library of approximately 36,000 small molecules from the compound screening library curated by the Chemical Biology Consortium Sweden (CBCS, SciLifeLab, details in supplementary materials). An overview to the screening workflow is outlined in Figure 3a. Controls included co-cultures exposed to DMSO as a vehicle control and <u>c</u>yclo<u>h</u>e<u>x</u>imide (CHX), a cytotoxic compound that blocks protein synthesis^46^ (Table 1, plate map in Supplementary figure 3b). Plates were then cultured for five days in presence of compound before analysis. In total, 74 measures including different morphological parameters such as covered area (of CAF subtypes and organoids, separately) and object intensities were used for multivariate analysis (Supplementary table 1). Data from these cultures were then processed and analysed both by multivariate approaches as well as a more traditional analysis focusing on fewer variables. Results from these analyses were used in conjunction to identify a shortlist of candidate compounds for further examination.

**Figure 3.**
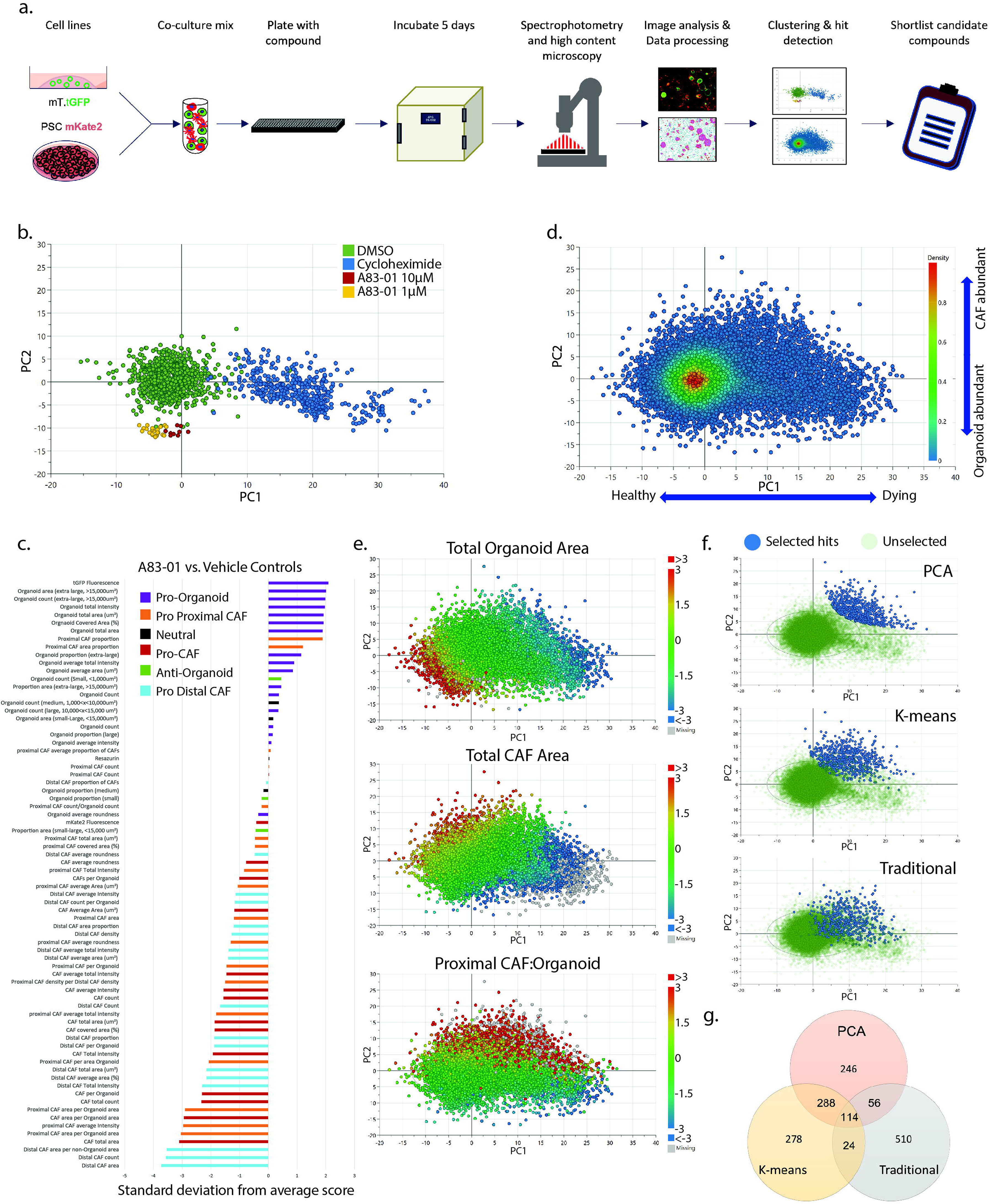
Drug screening and hit selection of a SciLife compound library. **(A)** Graphical workflow of cell and compound handling for screen set up. <u>M</u>urine pancreatic ductal tumor organoids (mT) and tumor naïve murine <u>p</u>ancreatic stellate <u>c</u>ells (PSCs) were each lentivirally transfected to stably constitutively express turbo-green fluorescent protein (tGFP) in the case of mT and mKate2 (a red fluorescent protein) in the case of PSCs. These cell lines, cultured separately, underwent a single cell dissociation, and mixed together at a ratio of 1:2 (1000 mT.tGFP cells: 2000 PSC.mKate2 cells per well) in matrigel and plated. Each well was then supplemented with media containing test compounds or control media and incubated for five days prior to analysis by spectrophotometry, microscopy by two imaging platforms and a resazurin end-point assay. These data were processed and analysed to identify candidate compounds with which to proceed for further analysis. **(B)** <u>P</u>rincipal <u>c</u>omponent <u>a</u>nalysis (PCA) of co-cultures treated with compound library together with control compounds. Each point represents the data from an individual well. Displayed are only the DMSO vehicle controls (green) and cycloheximide treated cultures (blue) as controls for normal co-culture growth and cytotoxic treated cultures respectively. Also displayed are example co-cultures treated with A83-01 at 1μM (yellow) and 10μM (red). Points are plotted on the first two principal components (R^2^= 0.365 and 0.176 for principal component one and two respectively). **(C)** A group to average comparison of the A83-01 treated co-culture wells compared to the average values from co-cultures in the initial screen modelled in the PCA. Displayed are the 74 metrics used in the co-culture model (y-axis) and the standard deviations away from the dataset average for those co-cultures treated with A83-01 as a whole (x-axis). The bars are pseudo-colored according to broad interpretation of what that metric associates with phenotypically. **(D)** A density plot of the PCA of all compounds tested in the initial screen using the same model as in Figure 3b. Each point represents one co-culture treated with a compound. Also indicated are simplified interpretations of the phenotypic features of the first and second principal components (blue arrows). **(E)** PCA of the treated compounds pseudo-colored according to their scaled values (Z-score) for total organoid area, total CAF area and the ratio of proximal CAFs: distal CAFs accounting for the area available to each (Data derived from the minimax analyses, variables #5, #12 and #70 from Supplementary table 1). **(F)** Visual representation of the top 704 hit compounds selected for further analysis by each selection method used including PCA, K-means and a traditional screening approach. Hits identified by each method are highlighted in blue, with unselected compounds in green. **(G)** A Venn diagram of hits as identified by the three different selection methods; PCA-based, K-means based and by a traditional approach.

**Table 1:** Media and culture composition for screening plates. Well composition within screening plate by treatment in terms of number of wells per plate, cell composition and media composition

| Name | Nº wells /plate | Nº Tumor cells | Nº PSCs | Final Media composition |
| --- | --- | --- | --- | --- |
| Treatment Co-Culture | 352 | 1,000 | 2,000 | Reduced media; 10.5 µM Y-27632; 10µM compound; 0.1% DMSO |
| Control Co-Culture | 8 | 1,000 | 2,000 | Reduced media; 10.5 µM Y-27632; 0.1% DMSO |
| Tumor Reduced mono-culture | 8 | 1,000 | 0 | Reduced media; 10.5 µM Y-27632; 0.1% DMSO |
| Cyclohexamide Co-Culture | 4 | 1,000 | 2,000 | Reduced media; 10.5 µM Y-27632; 1 µM Cycloheximide; 0.1% DMSO |
| Tumor Feed mono-culture | 4 | 1,000 | 0 | Feed Media; 10.5 µM Y-27632; 0.1% DMSO |
| PSC Reduced mono-culture | 4 | 0 | 2,000 | Reduced media; 10.5 µM Y-27632; 0.1% DMSO |
| Mgel reduced background | 4 | 0 | 0 | Reduced media; 10.5 µM Y-27632; 0.1% DMSO |

Hit selection was conducted in two successive rounds of screening. An initial screen on the compound library of ∼36,000 compounds was performed on one biological co-culture replicate at a concentration of 10μM per test compound. From these, 704 compounds were shortlisted and re-screened at three concentrations (5, 10 and 20μM) in the same biological replicate. False positive hits were identified and discarded from further consideration, and only those compounds that were effective over the concentration range were considered potential candidate compounds.

### Results of Initial Screen

When screen-wide data from the initial screen of co-cultures treated with test compounds, DMSO vehicle controls and CHX were analysed together by principal component analysis (PCA), the differences between the DMSO vehicle controls and CHX treated co-cultures were apparent, forming separate clusters that were distinct primarily along <u>p</u>rincipal <u>c</u>omponent (PC) one (Figure 3b). Furthermore, co-cultures that were treated with A83-01 also clustered distinctly from DMSO vehicle controls along PC2 and also demonstrated a dose-dependent separation in clustering along PC1 (10μM compared to 1μM). Whilst CHX treated co-cultures demonstrated very little cell growth as expected, A83-01 treated co-cultures demonstrated greater organoid growth, with a concomitant reduction in proximal CAF formation around these organoids, indicating also a reduction in myCAF formation (Figure 3c, Figure1d,I,j). In addition, a comparison of those cultures treated with A83-01 compared with the average values for co-cultures in vehicle control reveals that A83-01 treatment results in high scores of variables correlating to larger organoids (purple) and variables associated with reductions in CAF phenotypic measures (red, orange, cyan Figure 3c), altogether demonstrating a phenotype where pharmacological intervention has increased tumor cell growth that is undesirable for any therapeutic translation.

When all treated co-cultures were plotted on the PCA, it was identified that the majority of cultures overlapped with the DMSO vehicle control, which was expected since it was anticipated that most compounds would not exert an effect on the co-cultures (Figure 3d). A closer inspection of the variable loadings for PC1 and PC2, that is, the measures that contributed to separation along these PCs, revealed that PC1 was primarily associated with variables describing culture viability, with considerable contributions from organoid size, the resazurin reduction capacity assay and bulk fluorescence measures for turboGFP and mKate2, which all associating negatively with PC1 (Figure 3d, Supplementary figure 4a). In contrast, loadings for PC2 were not related to the overall metabolic activity of the cultures, since for example the resazurin assay measures contribute little to this component and instead PC2 seems to be related to the cell composition of the cultures (Figure 3d, Supplementary figure 4b). In general, cultures positive for CAF related metrics were positive on PC2, whereas cultures with more turboGFP and larger organoids associated negatively on PC2.

Altogether, this indicated that the bottom left quadrant of the PCA associates with cultures that displayed prominent tumor organoid growth and upon examination of this quadrant, these cultures indeed had much larger organoids than average (Figure 3e, top), without severe inhibition of overall CAF growth (Figure 3e, middle; Supplementary figure 4c). Although there was an increase in the proportion of proximal CAFs of all CAFs, likely due to the increased area of opportunity for proximal CAF detection around a larger area of organoids, the area of proximal CAFs per area of organoids was much reduced which is similar to the effect seen with A83-01 supplemented co-cultures (Figure 3c,e; Supplementary Figure 4c). Conversely, the top right quadrant was composed of cultures that had the opposite phenotype, such as reduced organoid growth and a CAF subtype balance that was shifted in favor of proximal CAFs compared to average (Supplementary figure 4d). Further, CAF growth was not especially reduced in this region, indicating that compounds here are disproportionately inhibiting tumor growth as opposed to CAFs (Figure 3e, middle). Altogether, this top right region appeared to represent an ideal phenotype for reducing tumor cell growth, without substantial inhibition to overall CAF growth but a shifted increase in proximal CAF per organoid. Therefore, candidate compounds for follow up study were selected from this region according to the product of their PC1 and PC2 coordinates (Figure 3f, top). These identified compounds likely alter the balance of CAF subtypes in favor of myCAF formation which in turn inhibit tumor organoid growth.

The PCA approach for compound selection utilized only the first two PCs of the dataset that together explain ∼54% of the total variance in the data (PC1 R^2^=0.365, PC2 R^2^=0.176). In total, 11 PCs were identified, cumulatively explaining 88.4% (R^2^) of the variance in the dataset (cumulative Q^2^= 0.804, Supplementary figure 5a). Given that approximately 35% of the explainable variance in the dataset was not considered in the PCA selection and that the PCA model was built by incorporating the data from the control cultures (DMSO vehicle and CHX) we explored utilizing an approach un-biased by data from controls. In this unbiased approach we analyzed the data from co-cultures treated with test compounds by considering only those co-cultures exposed to test compounds.

The overall idea was to allow the various phenotypes of these co-cultures to be examined without influence in the modelling from control cultures. The aim was to remove noisy variance in the data by subjecting it to another PCA, acquiring as many PCs that had explanatory power over the variance in the dataset as determined by the cumulative Q^2^ value calculated from the cross-validation process of model generation. Once noise in the data is removed and a dimensionality reduction performed by converting the initial 74 metrics into fewer PCs with explanatory power, the loadings for the PCs could be used for a K-means clustering algorithm to explore the relationship between culture clustering across these PCs in n-dimensional space. Once clusters are identified, they could be examined with respect to their members’ z-scores for the initial 74 metrics for each co-culture, from which a phenotype could be interpreted.

In this approach, the raw data of the 74 metrics from the initial screen were again subject to a non-stringent outlier removal and plate-wise scaling, however only considering those data from co-cultures exposed to test compounds, and not including any influence from controls. In total, 13 PCs were identified, cumulatively explaining R^2^= 90.8% of the variance in the dataset (cumulative Q^2^= 0.804, Supplementary figure 5b). We explored this data with a K-means clustering analysis, using the scores from the 13 PCA principal components as an initial dimensionality reduction step. The scores for each PC were extracted from each co-culture and re-scaled to give each PC equivalent weighting before being subject to a k-means clustering algorithm. The data were examined by the gap statistic algorithm to identify the ideal number of clusters for k-means clustering, (Supplementary figure 5c). Ultimately, 14 clusters were used for the K-means analysis (Supplementary figure 5d). Each cluster was then examined in terms of its members’ z-score for the 74 initial measures determined from each co-culture which were used to determine the phenotype captured by each cluster. In particular, cluster 14 demonstrated a phenotype of reduced organoid growth with altered, but not overall reduced, CAF metrics (Supplementary figure 5e). This cluster contained 835 co-cultures. To select the top 704 as candidates for further analysis, we selected cultures that were furthest from the origin of all 13 of the PCs used in the analysis, reasoning that the origin of a PCA definitionally represents the mean variance of all data modelled and consequently the most “normal” (especially given the large number of compounds that did not affect co-culture growth (Figure 3d). Given that we are searching for compounds that yield unusual phenotypes, we excluded those 131 co-cultures in cluster 14 that were closest to the origin of these 13 PCs. An overview of how those 704 compounds selected from cluster 14 appeared in the original PCA are highlighted in Figure 3f, middle.

Finally, a simultaneous analysis was performed using a more traditional screening approach that employs fewer measures. Specifically, four measures were used for the analysis (Table 2). The aim of this analysis was to select compounds that reduced mT growth, without showing signs of being generally cytotoxic. In this case, compounds were considered generally cytotoxic if they reduced the number of CAFs below that of the PSC mono-culture controls, since they generally grow very little if not stimulated in co-culture (Figure 1l). Thus, a reduction in PSC growth compared the PSC mono-cultures would indicate the compound being detrimental to PSCs, which is an undesirable outcome considering that PSCs are resident in healthy pancreas^47,48^. Alternatively, if the number of detected organoids was less than 50% of those in the DMSO vehicle controls, the compounds were deemed too potent and excluded from further analysis. Likewise, compounds must reduce the overall mT growth as determined by size, without increasing the total number of organoids. Further, “extra-large” organoids (>15,000μm^2^) should not contribute to the majority of overall organoid area, which would indicate an escape mechanism from the activity of the compounds. Compounds were ultimately rank ordered according to the total mT area in these cultures (Table 2, Supplementary table 1 metric #34) and the top 704 taken forward as candidate compounds from this method of selection, whose comparative position in the PCA plot are highlighted in Figure 3f, bottom.

**Table 2:**
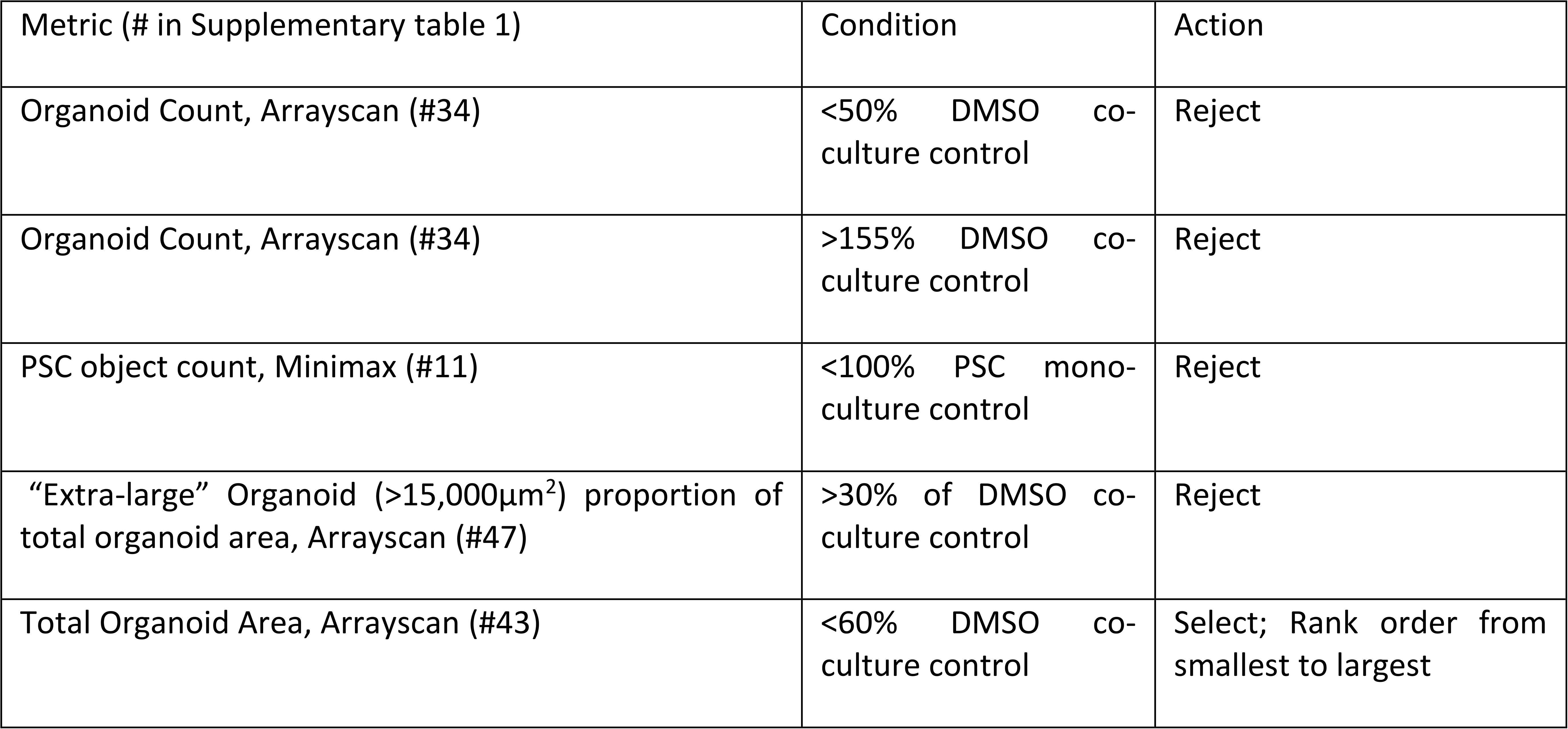
Traditional selection criteria from initial screen:

| Metric (# in Supplementary table 1) | Condition | Action |
| --- | --- | --- |
| Organoid Count, Arrayscan (#34) | <50% DMSO co-culture control | Reject |
| Organoid Count, Arrayscan (#34) | >155% DMSO co-culture control | Reject |
| PSC object count, Minimax (#11) | <100% PSC mono-culture control | Reject |
| “Extra-large” Organoid (>15,000µm<sup>2</sup>) proportion of total organoid area, Arrayscan (#47) | >30% of DMSO co-culture control | Reject |
| Total Organoid Area, Arrayscan (#43) | <60% DMSO co-culture control | Select; Rank order from smallest to largest |

Overall, the top 704 compounds selected from each method were compared together (Figure 3f, g). In total, 1,516 unique compounds were identified where 114 were identified in all three methods, and 368 were identified by any two methods (Figure 3g). This left 1,034 compounds that were uniquely identified by only one method each. To proceed to the hit verification phase, all compounds that were identified by multiple methods (n=482) were selected for further analysis, reasoning that the concordance between methods indicates a higher likelihood of a true hit. The remaining compounds were selected evenly from the remaining top ranked compounds from the three individual selection methods. Overall, the three selection methods are selecting compounds in a similar area of the PCA, where total organoid growth is low, and there is a high proximal CAF to organoid area.

### Hit verification screen

The 704 compounds identified from the initial screen were re-tested in the same screening setup (Figure 3a), and tested at three concentrations (5, 10, 20μM). To validate the multivariate analysis from the initial screen, a predictive <u>o</u>rthogonal <u>p</u>rojections to latent structures (OPLS) modelling approach was utilized. Here, the data from the initial screen and the new data from the hit verification screen were to be compared to validate that these compounds do behave as identified by the initial PCA approach. However, this required an alternative standardization method than the plate scaling approach used in the initial screen since the compounds for the hit verification screen are selected to deliberately skew the data distribution of the metrics examined. Therefore, data from the initial screen and data from the hit verification screen were plate-wise standardized according to each metric’s fold change of the mean of the DMSO vehicle control wells for each plate. When the data from the initial screen are re-examined by PCA, it’s seen that the loadings for PC1 (R^2^=0.98) and PC2 (R^2^=0.96) correlate very strongly between the two standardization methods, namely plate-wise scaling, and fold change of vehicle control (Supplementary figure 6a). This indicates that the datasets are robust to data standardization approach and that regardless of these two standardization methods, PC1 and PC2 are each still describing similar relationships.

To verify that compounds demonstrated the effects as observed in the initial screen, a predictive OPLS model was generated using a training dataset comprised of data from the DMSO vehicle controls (n=945), CHX treated co-cultures (n=447) and the 704 selected compounds from the initial screen (n=704, Figure 4a). The OPLS model had an R^2^ of 0.783 (X, modelling of metrics), 0.844 (Y, modelling of culture identity) and a Q^2^ of 0.839 (Supplementary figure 6b), with AUCs >0.995 for all three culture identities indicating that the data fits the model well, with good predictive capacity (Receiver operating characteristic, Supplementary figure 6c). Next, the new data from the hit verification screen was modelled in the OPLS model as test data for the model to predict whether the compounds demonstrated a phenotype that more closely resembled co-cultures treated with DMSO vehicle control, CHX or those data of the 704 compounds selected from the initial screen (Figure 4b). The majority of compounds tested clustered in the region that resembles data from the selected hits from the initial screen, even at 5μM (Figure 4b, top). There is a clear overall dose-response in the cultures, where the compounds at 5μM concentration are shifted closer to the DMSO vehicle control treated condition (Figure 4b, top), the 10μM condition leads co-cultures to overall present more like cultures from the initial screen (Figure 4b, middle) and at 20μM, cultures are shifted on average closer towards the CHX treated conditions (Figure 4b, bottom).

**Figure 4.**
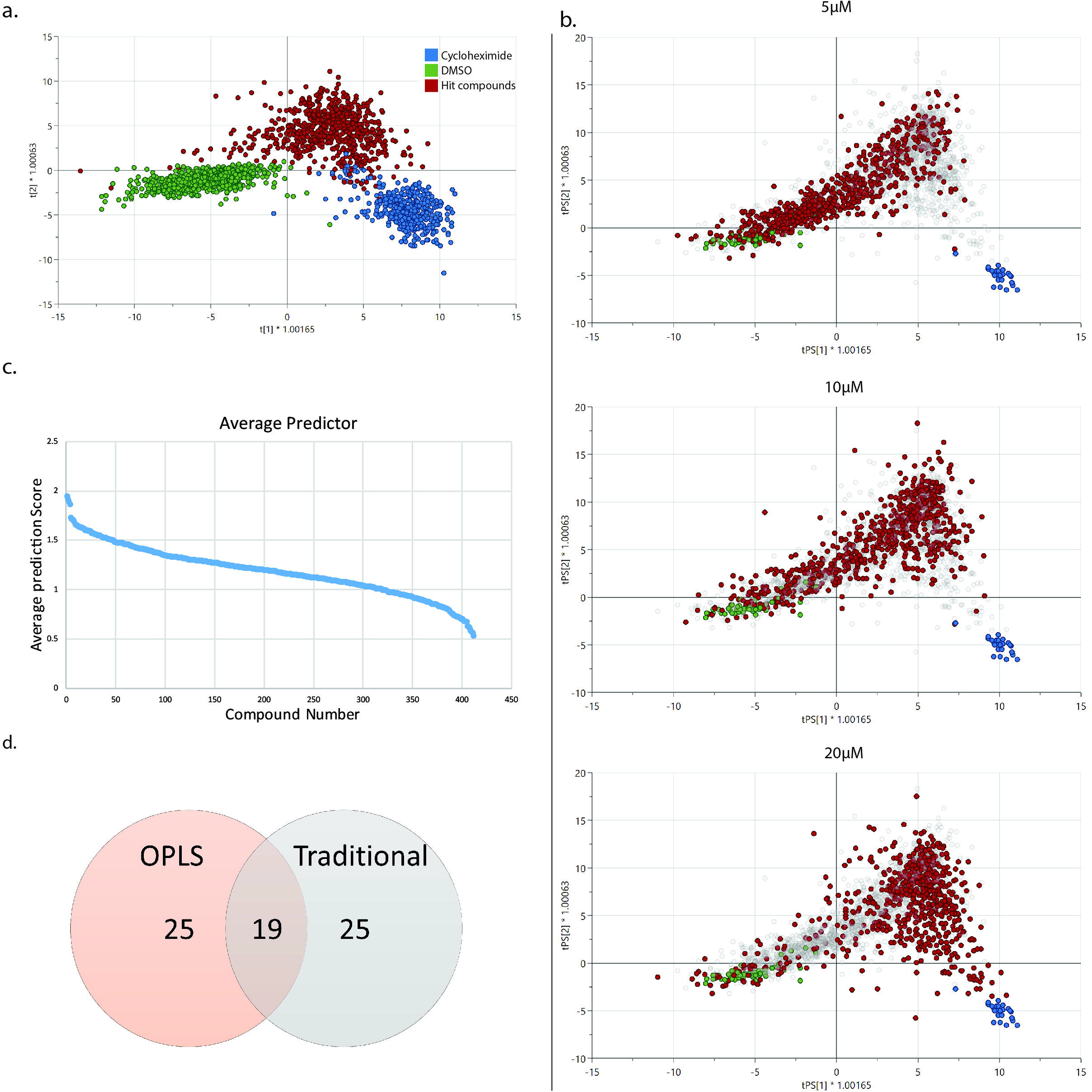
Hit verification screen analysis and selection. **(A)** Orthogonal projection to latent structures (OPLS) plot for training dataset from the initial screen. Each point represents the data from an individual co-culture. Depicted are the scores for the DMSO treated vehicle controls (green), cycloheximide treated co-cultures (blue) and those co-cultures of the 704 compounds selected from the initial screen (red). **(B)** OPLS plots of the hit verification screen modelled as a test dataset in the OPLS model. Each point represents the data from an individual co-culture. Depicted are the scores of the hit verification data and are colored according to their true identity as DMSO vehicle control (green), cycloheximide treated co-cultures (blue) and those co-cultures of the 704 compounds tested in the hit verification screen (red and grey). In this screen, three concentrations of compound were used: 5 (i), 10 (ii) and 20 (iii) μM. All data is shown in each graph, data highlighted in red are of different concentrations, where from top to bottom: 5μM, 10μM and 20μM. **(C)** Waterfall plot of remaining 412 compounds predicted to be a test compound at all concentrations by the OPLS model. Compounds are ranked by their average prediction score for being a test compound across the three tested concentrations of 5, 10 and 20μM. The Y-axis shows the average score and each integer on the x-axis refers to a different compound. Compounds are ranked from highest average score to lowest average score. **(D)** Venn diagram of the top 44 hits identified by the OPLS model and traditional method of hit verification. Of these, 19 hits were identified by both methods.

Compounds that resulted in cultures predicted to be either cycloheximide treated or resembled the DMSO vehicle control at any concentration were excluded from further consideration, leaving 412 compounds (58.5%). To select the top compounds for further consideration, two concurrent selection processes were undertaken; one by means of the OPLS predictive score as a test compound, and the other via a more traditional selection process similar to that used in the initial screen (Table 3). In the case of the OPLS selection, the remaining compounds were rank ordered according to their average predicted scores as likely test compounds across all three tested concentrations and 44 compounds could be selected (Figure 4c). Concurrently, the more traditional approach was based on fewer variables with an emphasis on selecting compounds that reduced organoid growth, without reducing overall PSC/CAF growth to avoid selecting for generally cytotoxic compounds (Table 3). The remaining compounds were rank ordered according to the total area of organoids measured by the Arrayscan (from low to high) in the 5μM condition for each compound and similarly 44 compounds were selected. Between the two approaches, 19 compounds were identified by both methods (Figure 4d). The remaining 25 compounds were selected evenly from the top ranked remaining compounds identified by each method.

**Table 3:** Traditional selection criteria from hit verification screen:

| Metric (# in Supplementary table 1) | Condition | Action |
| --- | --- | --- |
| Total number PSC, Arrayscan (#48) | <50% DMSO co-culture control at any two concentrations | Reject |
| PSC Object count, Minimax (#11) | <85% PSC mono-culture control at any concentration | Reject |
| Resazurin, Spectramax I3x (#3) | <20% DMSO co-culture control at any concentration | Reject |
| “Extra-large” Organoid (>15,000µm<sup>2</sup>) proportion of total organoid area, Arrayscan (#47) | <40% DMSO co-culture control at 10 and 20 µM | Select |
| Total Organoid Area, Arrayscan (#43) | <37% DMSO co-culture control at any concentration | Select; Rank order by 5µM condition |

### Identification of GNF-5 & Dose Response

The top 44 compounds identified from the hit verification screen were cross-referenced with the CBCS screening library annotations. One of the compounds was identified as GNF-5, an allosteric inhibitor of the Bcr-Abl fusion protein^45^. The ABL1 proto-oncogene is expressed in many tissues and is found in many cancer types, including pancreatic cancer^49^ (Supplementary figure 7a). ABL1 is expressed in pancreatic ductal cells and fibroblasts and is linked to ECM organization^50^ (Supplementary figure 7b). A search on functional protein association networks revealed ABL1 clusters together with ACTA2 (α-SMA), ACTA1, CTNNB1, APC and BCL9 pointing to a relevant molecular mechanism in the PDAC tumor-stroma context (Supplementary figure 7c). Given GNF-5’s use as a therapeutic against ABL1, we investigated its effect on tumor growth in co-cultures. Dose-response assays were conducted in three different biological replicates at increasing concentrations of GNF5. The effect in co-cultures was compared versus mono-cultures of tumor organoids and mono-cultures of non-tumor murine pancreatic ductal epithelial (“normal”, mN) organoids derived from healthy pancreas. Here, the aim was to explore whether there was a therapeutic window between tumor and non-tumor pancreatic ductal organoids, indicating that it would not be generally cytotoxic against pancreatic epithelial cells. Displacement of the mT.tGFP-growth curves in non-tumor monocultures compared to the PDAC co-cultures suggests GNF-5 has a therapeutic window (Figure 5b).

**Figure 5.**
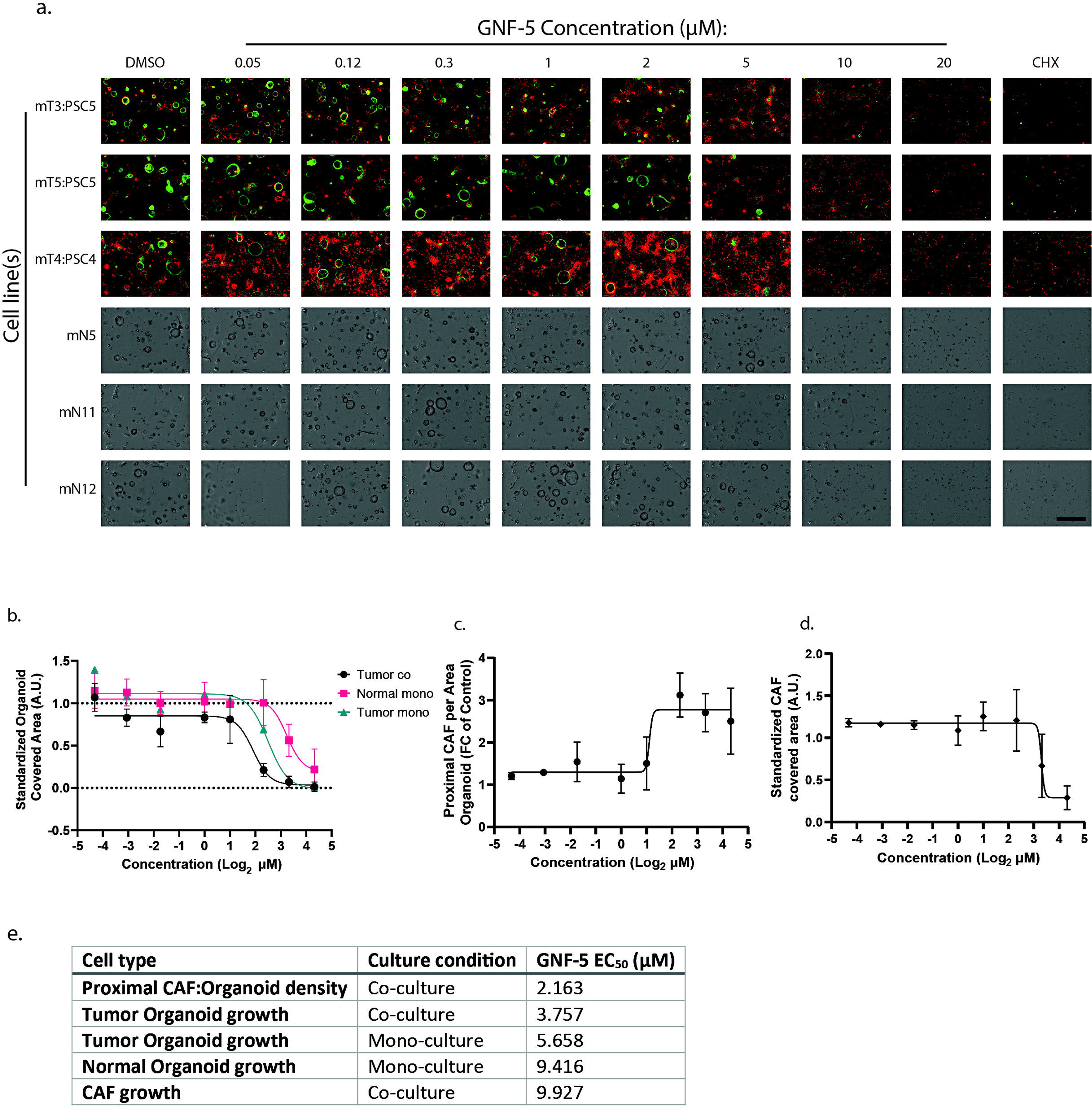
GNF-5 dose-response in normal and tumor pancreatic organoids. **(A)** Compilation of micrographs of cultures depicting GNF-5 dose-response together with representative images of those cultures treated with DMSO vehicle control or cycloheximide. Compound concentration alters by column, and different rows represent the different biological replicates. Shown are representative composite fluorescence images and brightfield images of the three different biological replicate tumor co-cultures and non-tumor mono-cultures, respectively. Micrographs were all taken at the same magnification and scale bar in bottom right micrograph represents 1,000μm. **(B-C)** Quantification of the dose responses of tumor co-cultures (n=3 biological replicate combinations), tumor mono-culture (n=1 biological replicate) and non-tumor mono-cultures (n=3 biological replicates) treated with GNF-5 for five days, for **(B)** organoid percentage covered area, **(C)** proximal CAF area per organoid area and **(D)** total CAF covered area without further subclassification (minimax). Data are standardized with respect to the mean DMSO vehicle controls and mean cycloheximide treated cultures on a scale of one to zero respectively, except for proximal CAF area per area Organoid, which is standardized as a fold change from the mean DMSO controls only. Data from C) and D) were collected from biological replicate (n=3) co-cultures. Error bars represent standard deviation. **(E)** Effective concentration 50 (EC_50_) values calculated for dose responses seen in (B-D), modelled by non-linear regression (GraphPad).

To quantitatively compare the cultures, the measures of organoid percentage well covered area after culture exposed to compound for five days were used. In the case of tumor organoid co-cultures, the measure of green fluorescence was used to determine the covered area, whereas in mono-cultures brightfield measures were used, since these lines did not express a fluorophore. An additional dose-response plate examined the growth of tumor mono-cultures (the same cell line that had been used in the initial and hit verification screens) which could be measured both in terms of its turboGFP fluorescence, as well as by brightfield microscopy. The two measurement approaches correlated reasonably well (brightfield vs. fluorescence measures of organoid covered area R^2^=0.733, Supplementary figure 8). The percentage covered area measures were standardized to allow comparison between cell lines. Therefore, a comparison on the relative dose-response of GNF-5 on the cultures with respect to organoid percentage covered area was performed and modelled by non-linear regression (Figure 5b). Tumor cells in co-culture were more susceptible to the effects of GNF-5 than non-tumor organoids, where the dose-response curves are significantly different and cannot be likely explained by fitting the same non-linear regression (p<0.0001, Comparison of fits, GraphPad, Figure 5b).

GNF-5 increased the proximal CAF per organoid area ratio in the same effective concentration range as against tumor organoids in co-culture (Figure 5c). Further, in the case of co-cultures, the effect of GNF-5 on the total CAF covered area, regardless of subtype (Figure 5d), indicated CAFs were less susceptible to the effects of GNF-5 than tumor cells. CAF growth was affected at similar concentrations (EC_50_=9.9 μM) to those of non-tumor organoid mono-cultures (EC_50_=9.4 μM) (p=0.300, Comparison of fits, GraphPad, Figure 5b,d,e). GNF-5 inhibited tumor cell growth in co-culture (EC_50_=3.8 μM), with an accumulation of proximal CAFs around tumor organoids (EC_50_=2.2 μM) at lower concentrations than when general cytotoxicity against CAFs and non-tumor organoids is demonstrated (Figure 5e). Furthermore, the dose responses indicated that tumor organoids in co-cultures are more susceptible to the effects of GNF-5 than tumor organoids in mono-culture (EC_50_=5.7 μM), which indicates an effect of GNF-5 via altered tumor-CAF interactions (Figure 5b-e). Taken together, GNF-5 emerges as a potent example of a compound that can potentiate the cancer-inhibiting properties of CAFs; a finding that illustrates the therapeutic strategy of harnessing the tumor-stroma to restrain cancer growth.

## Discussion

In this study we show that the balance between iCAFs and myCAFs in the co-culture model can be manipulated by chemical intervention which can inhibit cancer cell growth. We start with a specific example of TGF-β pathway manipulation using A83-01 and recombinant TGF-β which demonstrated pronounced effects on myCAF and iCAF formation which in turn associated with altered tumor organoid growth (Figures 1, 2). In particular, an increase in myCAF density around tumor organoids was concomitant with reduced cancer cell growth, indicating that in at least this context, shifting the CAF subtype balance in favor of myCAFs restricts cancer cell growth (Figures 1, 2). Consequently, for therapeutic application, we sought to identify novel compounds that alter the balance of CAF subtypes in co-culture and reduce tumor growth. Analysis focused on detecting changes in the symbiotic relationship between the tumor cells and stromal cells, reasoning that these compounds may affect tumor progression *in vivo*.

The identified mechanism of increasing the myCAF to iCAF ratio to inhibit tumor cell growth represents just one potential therapeutic avenue of CAF subtype manipulation. Knowledge of all the CAF subtypes in the co-culture model remains incomplete, and several other CAF subtypes are being discovered both in the context of PDAC^43,51–54^ but also CAF functional heterogeneity is being discovered in other cancer contexts such as in breast^44,55–57^, oral^58^ and colorectal cancers^59^. The presence of novel CAF subtypes described in PDAC, including <u>a</u>ntigen-<u>p</u>resenting CAFs (apCAFs) and <u>me</u>tabolically active CAFs (meCAFs) with proposed pathophysiological roles in the disease, has not been interrogated in the co-culture model^53,54^. Furthermore, heterogeneity of normal fibroblasts within the pancreas has recently been described^60,61^. These distinct normal fibroblast progenitor populations have been demonstrated to differentially contribute to functionally and phenotypically distinct CAF subtypes in PDAC. Both iCAFs and myCAFs are no longer considered as homogeneous populations but rather are further divisible into subpopulations governed by the normal fibroblast progenitors from which they arise. Depending on their cellular origin iCAFs and myCAFs may orchestrate variable influences on tumor cells and disease progression^61^. The characterization of PSCs used in the screen in the context of normal fibroblast heterogeneity is therefore of importance to better understanding the CAF composition of the co-culture model system. The relationships of these novel subtypes to the tumor cells and each other still need to be fully elucidated, however this ever increasingly complex network yields additional potential avenues for therapeutic TME manipulation. Thus, the co-culture model is heterogeneous in cell composition; a fact that is compounded by multiple feedback loops between cell types and the ability of CAF subtypes to interconvert. In such a heterogeneous system, it is difficult to envision *a. priori* what specific effect(s) a novel therapeutic may have on co-culture composition, particularly by imaging methods, indicating that the effects of novel therapeutics may be difficult to interpret in a univariate approach. Furthermore, the heterogeneous nature of the co-culture model results in cultures that are themselves varied, which is not well suited for the development of robust Z-prime scores that are the mainstay of more usual traditional screening methods.

To overcome both problems of culture variance and limitation of a comprehensive univariate measure, we employed multiple approaches to analyze a co-culture model of PDAC including multivariate modelling and more traditional analyses incorporating fewer variables. With respect to the PCA multivariate analysis, the objective was to first model the normal pattern of co-culture growth in this model system (DMSO vehicle control, Figure 3b). Then, in combination with a set of control co-cultures exposed to the cytostatic compound CHX, we were able to anchor our analysis in terms of overall growth by PC1. Finally, by examination of another condition known to alter the phenotype of interest (by addition of A83-01), we were able to identify different phenotypes within our PCA, and in particular with respect to PC2. A similar multivariate approach aimed to allow the measures from the co-cultures treated with compound cluster the co-cultures directly into those with similar phenotypes. By first performing a PCA analysis on these data, variance in the dataset without explanatory power is omitted as well as simultaneously undergoing a dimensionality reduction step. K-means clustering of the compounds with respect to their PC scores revealed 14 clusters, the members of which were analysed with respect to initial measures of the co-cultures. Finally, a more traditional approach focusing on few variables was used to identify potential compounds of interest.

For all the selection approaches, there were several key phenotypic criteria required for identifying promising compounds with potential therapeutic application. Firstly, the compound should act to produce an abnormal growth pattern in co-culture outside the range of the control co-cultures. Secondly, the potential therapeutic should act to reduce tumor organoid growth. Finally, it should not act in a cytotoxic manner with respect to the CAFs in the culture. These criteria allow for the identification of compounds that may either act on the tumor cells directly or by interfering with the relationship between the cell types present in the culture. Overall, the three approaches identified compounds in a similar region of the PCA space (Figure 3f). Of the top 704 compounds identified by each method, 114 were identified by all three methods, which is extremely unlikely if selections were made at random, indicating strong concordance between the methods.

These selected compounds were tested in a hit verification screen to check for reproducible activity so that false positive compounds could be omitted from further consideration. To make this comparison new data from the hit verification screen were tested against an OPLS model trained on the data from the initial screen. True positive compounds ought to demonstrate similar effects on co-culture as seen in the initial screen. Indeed, most compounds (58.5%) appeared to behave similarly to those selected from the initial screen without demonstrating cytotoxicity equivalent to CHX nor a lack of potency at any tested concentration. Selection here was performed by selecting those compounds that were predicted to be most like the hit compounds from the initial screen across all concentrations. That most compounds appeared to be true hits was also observed when examining the compounds by the more traditional approach. Selection of compounds was consequently made more stringent (Table 3). The top 44 compounds determined by each method were compared where 19 compounds were identified in the top 44 by both methods which is, again, unlikely.

This shortlist of 44 compounds uncovered GNF-5 as a potent inhibitor of tumor cell growth in co-culture that appeared to also be less potent against non-tumor organoid and tumor organoids in mono-culture, and CAF growth (Figure 5). One important caveat to consider is that the non-tumor organoids cannot be cultured in the reduced media conditions of the tumor co-culture, and were maintained in complete organoid feeding medium, which may have contributed to improved tolerance of non-tumor organoids to GNF-5. However, the effect of GNF-5 on CAF growth in co-culture mirrored that of the non-tumor organoids, supporting the idea that GNF-5 is disproportionately inhibiting the growth of the tumor cells and not generally cytotoxic to pancreatic ductal epithelial cells. Further, the addition of GNF-5 increases the proportion of proximal CAFs to organoid area, indicating that this effect may be mediated via CAF subtype manipulation.

GNF-5 is an inhibitor of the ABL1 proto-oncogene, which encodes a protein tyrosine kinase involved in cell division, adhesion, differentiation and stress response^50,62^. It is overexpressed in different cancer types including lung adenocarcinoma and <u>c</u>hronic <u>m</u>yeloid leukemia (CML), where therapeutic ABL kinase inhibitors sensitize the tumor to chemotherapy by promoting tumor cell differentiation and apoptosis in metastatic cancer cells^63–66^.

Here, we demonstrate that GNF5 can also be used to modulate the tumor-stroma environment with direct implications for tumor growth since its effect against tumor organoids in co-culture was more potent than seen in tumor organoid mono-cultures (Figure 5). This also has potential implications with respect to chemosensitivity since abundance of CAFs has been previously linked to chemoresistance^67^. Indeed, ABL1 is enriched in different cell types including pancreatic ductal cells and fibroblasts (Supplementary figure 7b). Single cell RNA-seq data from the human protein atlas indicates co-expression of transforming growth factor beta 1 induced transcript 1 (TGFB1I1), fibroblast growth factor (FGF)-10, growth differentiation factor (GDF)-7 and collagen 12 alpha 1 chain (COL12A1) in fibroblasts (Supplementary Figure 9). Which together suggests a role of ABL1 in myofibroblast formation besides integrin-mediated actin cytoskeletal remodeling and ECM organization (Supplementary figure 9).

Given the role of ABL1 in the pathogenesis of pancreas-related disease and its therapeutic potential in cancers driven by loss of function point mutations in TP53 and activating mutations in KRAS, it is plausible that ABL1 inhibition will also benefit pancreatic cancer patients^68^. However, ABL1 inhibitors with the exemption of imatinib have been shown to trigger vascular toxicity *in vivo*, prompting further research to explore combinational treatments to mitigate these unwanted effects^69^. Since increased Rho-associated kinase (ROCK) activity has been found to contribute to vascular toxicity in patients with CML after treatment with ABL tyrosine kinase inhibitors, we hypothesize that combinatorial treatments with ROCK inhibitors will reduce the risk of cardiovascular side-effects^45,70^. Remarkably, GNF5 combinatorial treatments together with first line chemotherapeutics such as nilotinib or docetaxel resulted in additive inhibition of BCR-ABL1 and increased survival ratios in BCR-ABL1-transduced bone marrow and lung cancer models confirming the GNF5 therapeutic potential^62^. Further preclinical assessment of GNF5 together with gemcitabine or 5-FU might therefore be of interest in the context of PDAC.

The co-culture model yields several experimental advantages over mono-culture models of PDAC. Firstly, the well-established interactions between tumor cells and CAFs in PDAC make the co-culture model a more biologically relevant setting for drug discovery in the PDAC context. Additionally, multiple targets are present within the screen through which a compound may act. For example, a compound may exert its effect by directly acting on any cell type directly or by modulating secreted signals between the tumor or CAF cell types. Conversely, compounds that affect tumor cells in mono-culture but are rendered inert by a rescue effect of the surrounding CAFs are removed from further consideration automatically in the screening process, thus avoiding investment in a compound that would later demonstrate limited clinical relevance. Further, PSCs are a cell type that is normally found in healthy pancreas, and so can act as an internal control for excluding compounds that are generally cytotoxic to these normal pancreatic cells.

In consideration of the cells used in this screen, pancreatic tumor organoids derived from the murine KPC model of pancreatic cancer were chosen since they contain defined mutations in KRAS and P53 that represent the condition found in the majority of PDAC patients. Identified compounds that act in this system are likely able to be generalized as a therapeutic. In contrast, using a patient-derived organoid cell line has the potential to identify compounds effective only at treating that specific patient, but not readily generalized to other cases. Of course, compounds effective in this murine *in vitro* model would still require validation in multiple contexts, including application to patient-derived PDAC organoids, ADMET studies as well as *in vivo* efficacy and safety testing for systemic effects. Further, uncovering the mechanism of action of such compounds requires consideration. Single cell transcriptomic analyses of treated co-cultures will allow enrichment signaling analysis, identification of genes involved in stromal interactions and drug response. High-throughput PISA assays in combination with proteomic approaches will also be key in determining drug-target interactions.

Overall, we have developed a screening approach to interrogate heterogeneous co-culture models. By multivariate means we have utilized the wealth of information afforded by high content image analysis to identify a novel anti-tumor role for GNF-5. GNF-5 demonstrates a specific window of activity where PDAC epithelial tumor cell growth is inhibited in co-culture but both PSC-derived progeny (CAFs) and non-tumor ductal epithelial organoids tolerate GNF-5 exposure. Further, GNF-5’s effects are associated with an altered CAF subtype formation in the co-culture promoting myCAF density around tumor organoids, indicating a cancer cell suppressive function of the myCAF subtype in this context. This finding underscores a role for the novel strategy of taking advantage of the inherent tumor restraining capacity of particular aspects of the tumor-stroma for the development and discovery of future therapeutics.

## Materials and Methods

### PSC Isolation and culture

PSCs were isolated from WT C57BL/6J mice as previously described^41^. Briefly, minced pancreata were digested at 37°C for 30 minutes in a dissociation buffer containing 0.05% collagenase P (Sigma-Aldrich) and 0.1% DNase I (Sigma-Aldrich) in <u>G</u>ey’s <u>b</u>alanced salt solution (GBSS; Sigma-Aldrich, G9779). Digested pancreata were then filtered through a 100-µm nylon mesh and washed in GBSS with 0.3% <u>b</u>ovine serum <u>a</u>lbumin (BSA) and 0.1% DNase I. Following centrifugation, the pellet was re-suspended in 9.5 ml GBSS with 0.3% BSA and 43.75% Histodenz (Sigma-Aldrich, D2158). Six ml GBSS with 0.3% BSA was layered on top of the cell suspension and the gradient centrifuged for 20 min at 1,400 RCF (break switched off). The cells in the band above the interface between the Histodenz and GBSS were harvested, washed in PBS, and plated. PSCs were cultured on tissue culture treated plastic with media composed of DMEM (Sigma-Aldrich, D5796) with 5% FBS (Gibco, 12657029) and penicillin/streptomycin (Gibco). PSCs were passaged using TrypLE express (Gibco, 12605) to dissociate for 2 min at 37°C, washed in DMEM supplemented with 5% FBS then centrifuged and re-suspended in culture media and finally re-plated at a 1:10 split ratio.

### Mouse organoid isolation and culture

The organoid lines were isolated from KPC mice with histologically verified PDAC and derived as previously described^71^. Briefly, tumor tissue was minced and digested at 37°C for 12 h in a dissociation buffer containing 0.012% (wt./vol.) collagenase XI (Sigma-Aldrich, C7657) and 0.012% (wt./vol.) dispase (Gibco, 17105041) in DMEM (Gibco, D5796) containing 1% FBS (Gibco). The tissue debris was allowed to settle, and the dissociated cells were pelleted and washed in Advanced DMEM/F12 (Gibco, 12634) and seeded in growth factor reduced Matrigel (Corning, 356231). Organoids were cultured in complete murine pancreatic organoid feeding media (Boj et al., 2015); Advanced DMEM/F12 supplemented with 1x GlutaMAX (Gibco, 35050), 1x Hepes (Gibco, 15630), 1x B27 (Invitrogen, 17504044), 1.25 mM N-Acetylcysteine (Sigma-Aldrich, A9165), 10 nM gastrin (Tocris, 3006), 50 ng/ml EGF (Thermofischer, PMG8041), 10% RSPO1-conditioned media, 100 ng/ml Noggin (PeproTech, 250-38), 100 ng/ml FGF10 (PeproTech, 100-26), and 10 mM Nicotinamide (Sigma-Aldrich, N0636).

To passage, organoids were washed out from the Matrigel using cold splitting media (Advanced DMEM/F12 supplemented with 1x GlutaMAX (Gibco) and 1x Hepes (Gibco)), and mechanically dissociated into small fragments using small aperture glass pipettes, and then seeded into fresh Matrigel. Passaging was performed at a 1:8 split ratio roughly twice per week. To create frozen stocks, organoids were passaged and mixed with Recovery Cell Culture Freezing Medium (Gibco, 12648010) and cryopreserved using standard procedures. Cultures were thawed using standard thawing procedures, washed once with splitting media, and seeded in Matrigel with organoid media supplemented with 10.5 µM ROCK Inhibitor Y-27632 (Sigma-Aldrich, Y0503) for the first passage.

### TGF-β Pathway modulation

Unless otherwise stated, cells were treated with 1µM of recombinant human A83-01 (TOCRIS), or 20ng/ml recombinant TGF-β (R&D Systems, 7666-MB-005) for 5-6 d before processing and analysis by described assays.

### Co-culture preparation for flow cytometry and transcriptomic assays

Co-cultures for flow cytometric or transcriptomic analysis were prepared by mechanical fragmentation dissociation in the case of organoids as described for their passaging. For PSCs, monolayer cultures were treated with TrypLE express for 2 min at 37°C and harvested in DMEM with 5% FBS, centrifuged and re-suspended in splitting media. Cultures were seeded into 24 well plates. Organoids were split in a 1:5 ratio and mixed with 15,000 PSCs per well in a volume of 50μL Matrigel/well. Mono-cultures were prepared identically but omitting the other cell type. Plates were incubated for 15 minutes at 37°C, 5% CO_2_ to allow the Matrigel to set prior to splitting media addition supplemented with relevant compound. Cultures were maintained for six days, with a media exchange on the third day after seeding.

### Co-culture processing for Flow Cytometry profiling of CAF subtypes

For intracellular flow cytometric staining, cultures were treated with GolgiPlug (BD, 555028) for 12 hours before harvesting cultures. Cultures were gently transferred into falcon tubes and dissociated by 2mg/ml dispase at 37°C for 20 minutes, followed by centrifugation and resuspension in a mix of TrypLE and DNase 10mg/ml at 37°C for 5 minutes then resuspended in DMEM/ 5% FBS to quench the TrypLE and cells were strained through a 70μm strainer. Suspensions were washed with splitting media, counted (Countess II, ThermoFisher) and normalized for equal cell number between samples. Then samples were resuspended in a Near Infra-red Live/Dead fixable stain (ThermoFisher, L10119) in PBS and incubated on ice for 15 minutes before centrifugation and wash steps in flow buffer (PBS, 0.2% Bovine serum albumin (Sigma-Aldrich, A4503), 2mM EDTA).

Samples were then fixed and permeabilized using a Cytofix/Cytoperm kit (BD, 555028) according to the manufacturer’s instructions. Cells were washed twice in perm/wash buffer (BD, 554723) then stained with the appropriate primary antibodies in Perm/wash buffer at 4°C for 30 min in the dark. Samples were stained with: conjugated Cy3-α-SMA mouse monoclonal antibody (Sigma-Aldrich, C6198), Anti-PDGFR-β antibody (Abcam, ab32570) and conjugated APC-anti-mouse-IL-6 antibody (Biologened, 504507). Samples were then washed twice in perm/wash buffer prior to staining with secondary antibodies and DAPI (Roche, 10236276001) in perm/wash buffer. Samples were stained with: Goat-anti-Rabbit IgG (H+L) Alexa Fluor 488 (Invitrogen, A32731). The stained populations were analyzed, with at least 20,000 events recorded per condition, using a BD FACSAria III flow cytometer and FlowJo software (Tree Star; Version 10).

### Co-culture processing for FACS and sequencing library preparation

Cultures were harvested by enzymatic single cell dissociation by 2mg/ml dispase at 37°C for 20 minutes, followed by centrifugation and resuspension in a mix of TrypLE and DNase at 37°C for 5 minutes. TrypLE was quenched by addition of ice-cold defined trypsin inhibitor (Gibco, R007100). Cells were strained through a 70μm strainer and resuspended in Live/Dead fixable stain (Invitrogen) in PBS and incubated on ice for 15 minutes before centrifugation and a wash in a solution of 0.1% bovine serum albumin in PBS. Samples were centrifuged and resuspended in 50uL PBS followed by addition of 3mL ice-cold methanol, gently mixed and incubated on ice for 10 minutes prior to centrifugation and resuspension in permeabilization buffer (EGTA, mgSO4, MOPS, NaCl, RNAsin, BSA, DTT, Saponin) on ice for 5 minutes.

Cells were centrifuged and resuspended in primary antibodies in permeabilization buffer on ice for 30 minutes. Primary antibodies used were anti-PDGFR-β (Abcam, ab32570) and TROMA-III (DSHB, AB_2133570). Samples were washed in permeabilization buffer the resuspended in secondary antibodies and DAPI in permeabilization buffer on ice for 20 minutes. Secondary antibodies used were Goat-anti-rat IgG (H+L) Alexa Fluor 647 (Invitrogen, A21247) and Goat-anti-Rabbit IgG (H+L) Alexa Fluor 488 (Invitrogen, A32731). Samples were then washed in permeabilization buffer and resuspended in MRDB (EGTA, MgSO4, MOPS, NaCl, BSA, DTT, RNAsin) and passed through a 70μM strainer and sorted by FACS using a BD FACSAria III flow cytometer for cells positive for TROMA-III (cancer cells) and cells positive for PDGFR-β (CAFs). For each cell type, 500 cells were sorted and collected directly in a lysis buffer of 0.2% Triton with RNAsin then immediately stored on dry ice and transferred to a minus 80°C freezer ready for cDNA library preparation.

For library preparation total RNA cell lysates were processed according to the previously established SMART-Seqv2 protocol^72^. Libraries fragment size distribution and concentration was assessed using Qubit dsDNA HS Assay Kit (Thermo Fischer Scientific, Q32851) and Agilent High Sensitivity DNA kit (Agilent Technologies, 5067-4626) on a bioanalyzer. Libraries were diluted to 6nM and pooled, 20pmol of pooled libraries were sequenced on Illumina NextSeq 500/550 High Output kit v2.5 (75 Cycles) (Illumina, 20024906).

### Transcriptomic Analyses

Illumina base call files were converted to FASTQ using the default Illumina bcl2fastq Conversion Software v2.20 pipeline with recommended parameters and sample file constructed with Illumina Experiment Manager. Adapter sequence corresponding to dT30, template switch oligo, as well as ISPCR sequence were removed using Cutadapt 3.1^73^.

Ribosomal RNA reads were identified for removal by mapping to 45S (NR_046233.2) and 5S (NR_030686.1) rRNA sequences using bowtie2.

Reads were mapped to the mouse reference genome primary assembly GRCm39 using STAR with default parameters. Duplicated reads were removed in a coverage-dependent manner using a custom script. Briefly, the coverage of exactly identical reads was reduced to that of adjacent sequence using a model based on the Poisson distribution and a cutoff of 0.05 for the probability of the observed coverage.

A custom annotation GTF file was generated from the Ensembl release 104 annotation by selecting transcripts with protein coding annotation. The sum of all exons through transcript variants was kept as a single transcript per gene. The read count per exon was derived with HTseq^74^ and the sum of count per gene was calculated as raw read count. Raw read count was processed for differential gene expression analysis using the DESeq2 package^75,76^.

Z-scores were calculated using the DESeq2 normalized read count. iCAF and myCAF signatures were derived from bulk sequencing data published by Öhlund, D. *et al*^41^. The composite genes of a gene signature were extracted from the normalized expression matrix and used to calculated Z-score for individual samples. A Z-score is a numerical quantification of the deviation from the mean in terms of standard deviations. Z-scores for each gene in a signature were first calculated as follows:

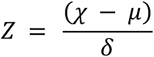

*χ* = normalized gene count

*μ* = mean gene count

*δ* = standard deviation

The mean Z-score of all genes in the gene signature was then taken to define a Z-score for a gene signature in an individual sample.

For analysis of transcription factor activity aligned bam files were uploaded for analysis using ISMARA software^77^

### Transfection of fluorophores and immortalization

Primary PSCs were immortalized by lentiviral transduction of the SV40 Large T-antigen. 293T cells were co-transfected with pBABE-puro-SV40-LT target plasmid (Addgene, Plasmid #13970) together with psPAX2 (Addgene, Plasmid #12260) and pMD2.G packaging plasmids (Addgene, Plasmid #12259) using X-treme GENE 9 DNA transfection reagent (Roche, XTG9-RO) to produce the lentiviral particles. Media was changed 24 h later, and lentivirus supernatant was collected after an additional 24 h. The supernatant was filtered through a 0.45-µm filter and aliquots were kept at –80°C. Primary PSCs were grown to 70% confluency and infected with the virus supernatant for 24 h. Virus was aspirated and fresh media containing puromycin for selection was added. To generate the lentiviral particles required for the fluorophore labelled cell lines, either pLVX-EF1α-turboGFP-WPRE-Neo (for turboGFP expression, Supplementary figure 2a; Vectorbuilder) or pLVX-EF1α-mKate2-WPRE-Neo (for mKate2 expression, Supplementary figure 2b; Vectorbuilder) target plasmid was transfected together with psPAX2 and pMD2.G packaging plasmids into 293T cells using lipofectamine 2000 transfection reagent (Invitrogen, 11668027). Media was changed 24h later and lentivirus supernatant was collected after an additional 24 h. The supernatant was filtered through a 0.45 µm filter and concentrated with Lenti-x concentrator (Takara Bio Inc., 631231) and 250 µl aliquots stored at −80°C.

To obtain turboGFP labelled organoids, an organoid culture from a single well of a 24 well plate, mechanically dissociated into small fragments using narrow aperture glass pipettes then digested briefly for 5 min at 37°C in TrypLE express (Gibco) which was then quenched using DMEM (Sigma-Aldrich) with 5% FBS (Gibco), centrifuged and re-suspended in a 250 µl aliquot of the pLVX-EF1α-turboGFP-WPRE-Neo lentivirus with 10 µg/ml polybrene infection/transfection reagent (Millipore-Sigma, TR-1003), transferred into a single well of a low attachment 48-well plate and centrifuged for 1 h at room temperature then incubated at 37°C 5% CO_2_ for 6 hours. The cell suspension was then collected in splitting media, centrifuged then re-suspended and seeded in Matrigel with organoid media supplemented with 10.5 µM ROCK inhibitor Y-27632 (Sigma-Aldrich) for the first passage. Two days later, media was exchanged with organoid media supplemented with 1mg/ml G418 (Roche, G418-RO) to begin selection.

To obtain mKate2 labelled PSCs, pLVX-EF1α-mKate2-WPRE-Neo virus supplemented with 10 µg/ml polybrene infection/transfection reagent (Sigma-Aldrich) was incubated with PSCs grown in monolayer for 24 h. Two days after infection, cells were treated with 1mg/ml G418 (Roche) for selection. Both the organoid and PSC cell lines were later enriched for highly fluorescent cells by fluorescence-<u>a</u>ctivated <u>c</u>ell sorting (FACS).

### Co-culture preparation for Proliferation Assays

For co-cultures, organoids and PSCs were enzymatically dissociated into single cell suspension. For PSCs, monolayer cultures were treated with TrypLE express for 2 min at 37°C and harvested in DMEM with 5% FBS, centrifuged and re-suspended in splitting media. For organoids, culture media was removed and cultures were disrupted by pipetting with 2 mg/ml dispase solution in splitting media, incubated at 37°C for 20 minutes, then harvested in splitting media and centrifuged. The pellet was re-suspended in TrypLE and incubated for 5 minutes at 37°C with continuous shaking then an equal volume of 2 mg/ml dispase solution added and the solution supplemented with DNase I then incubated for a following 10 min at 37°C with continuous shaking. Cells were then washed with splitting media centrifuged then re-suspended in splitting media and maintained on ice.

Unless otherwise stated, proliferation assays were performed in a 384 well plate format (Falcon 353962) with an initial seeding of 1,000 mT.tGFP and 2,000 PSC.mKate2 cells in a 15μL volume per well. Therefore, the single cell suspensions were counted (Countess II, ThermoFisher) and diluted to a concentration of 5.3×10^5^ PSC.mKate2 cells/ml and 2.7×10^5^ mT.tGFP cells/ml in splitting media. The resulting mixtures were mixed at a 1:1 ratio and finally mixed with an equal volume of Matrigel to give a final mixture that was 1:1 Matrigel: co-culture cell suspension sufficient for plating the required number of wells. The cell mixtures were then plated on ice, incubated at room temperature for 20 minutes, then incubated at 37°C for 30 minutes and then the desired culture media was added supplemented with 10.5 µM Y-27632 (Sigma-Aldrich).

For mono-cultures the process was identical except for the omission of the other cell type, resulting in initial seeding of either 1,000 mT.tGFP or 2,000 PSC.mKate2 cells per well in a 15uL volume of 1:1 Matrigel: mono-culture cell suspension in splitting media. Unless stated otherwise, cultures in proliferation assays were maintained in reduced media conditions; Advanced DMEM/F12 supplemented with 1x GlutaMAX (Gibco), 1x Hepes (Gibco), 10.5μM Y-27632 (Sigma-Aldrich).

Resazurin assays were conducted by using PrestoBlue Cell viability reagent (Invitrogen, A1326). For a culture’s end-point readout, 10% of the total volume of well contents (matrigel volume + media volume) at time of seeding were added and mixed by gentle shaking. Plates were incubated for one hour at 37°C before being read by spectrophotometry in the Spectramax i3x platform (excitation 550nm, bandwidth 15nm; emission 590nm, bandwidth 15nm; 6 flashes per read from bottom).

### Image processing for figures

All analyses on micrographs were performed on the raw image file or were processed as describe in the materials and methods section. However, for presentation in figures, micrographs were processed for better viewing as follows. Fluorescent image data from the minimax plate reader were subject to a background subtraction in Fiji^78^, converted to RGB image and brightness levels altered to improve visual dynamic range. For images from the 541nm channel, the minimum was 5, maximum 120. For images taken from the 713nm channel, images the minimum was 1, maximum 25. These images were then stacked into a composite using the merge channels function in Fiji, then converted and saved as an RGB image. Images from the brightfield channel were unaltered.

### High Throughput Screen

Compounds screened were acquired from the <u>C</u>hemical <u>B</u>iology <u>C</u>onsortium <u>S</u>weden (CBCS SciLifeLab, KI) screening library of approximately 36,000 compounds. Compounds were maintained under their storage conditions and transported to our labs ready for screening where they were maintained in DMSO at approximately −20°C prior to use in the screen (details in supplementary material “Description of CBCS screen Library”).

Co-cultures for the screen were prepared using the co-culture preparation described above using the lentivirally modified cell lines described above. The plate layout was designed to include 352 co-cultures treated with test compounds (10μM initial screen; 5, 10 and 20μM hit verification screen), eight control co-cultures treated with DMSO as a vehicle control, four control co-cultures with cycloheximide (250ng/ml, Sigma-Aldrich, C4859) added as a cytotoxic control, eight wells of tumor organoids in mono-culture in reduced media, four wells of tumor organoids in mono-culture in organoid feeding media, four wells of PSCs in mono-culture in reduced media conditions and finally four wells of seeded matrigel without cells in reduced media conditions (Supplementary figure 3b, Table 1). Cultures were seeded in the cell culture plate chilled on ice in 15 µl 1:1 Matrigel/splitting media mix per well according to the plate map (Supplementary figure 3b). When a plate was fully seeded, it was briefly spun at 200 RPM at 4°C for 15 seconds then incubated to allow matrigel to set as described above. Following incubation, 48 μl of relevant media was seeded for a final volume of 63 µl per well (Table 1). All conditions had a final concentration of DMSO (0.1% in the initial screen, 0.2% in the hit verification screen) and 10.5 µM Y-27632 (Sigma-Aldrich). Co-cultures were incubated at 37°C with 5% CO_2_ for five days prior to assay readout.

Plates were assayed sequentially by scanned well spectrophotometry of total fluorescence using the SpectraMax I3x platform (Molecular Devices) for a readout on turboGFP (read settings: Excitation: 485nm (bandwidth 9nm), Emission: 510nm (bandwidth 15nm) and mKate2 (read settings: Excitation 588nm (bandwidth 9nm); Emission 633nm (bandwidth 15nm)) fluorophores over 9 points per well too account for spatial heterogeneity in co-culture distribution throughout the well. Plates were then imaged using the Minimax function of the SpectraMax I3X with one field per well in the channels for transmitted light (6ms exposure), as well as the green (541nm, 10ms exposure) and red (713nm, 1000ms exposure) fluorescent channels. The SpectraMax stage was heated to 37°C throughout. Images collected by the Minimax imaging procedure were analyzed by the associated SoftmaxPro 7.0 software (Molecular Devices) in the green channel for organoid measures and also the red channel for PSC/CAF measures. CAF subtype classification for red channel images from the Minimax was performed using the machine-learning classification algorithm in SoftmaxPro 7.0 software where training for classification was supervised by manual review. All cultures in the screen were classified by the same version of the classification algorithm. The algorithms used for mask detection (both organoid and PSC/CAF) and cell classification (in the case of CAFs), which was trained in advance of the screen, can be found in the online supplementary materials

Plates were also imaged using the Arrayscan VTI coupled with live cell module (Thermo Scientific) high content imaging platform at 5x magnification over four fields per well. The Arrayscan stage was heated to 37°C with 5% CO_2_ throughout the imaging procedure. Images were acquired in two channels, one at excitation of 485nm (band 20nm) for an exposure time of 0.012 seconds per field and the other at excitation of 549nm (band 15nm) for an exposure time of 0.075 seconds per field. A standard laser autofocusing procedure was used using the 485nm channel on the first field per well which was then applied to the remaining three fields per well. Images from the 485nm channel were processed for background removal (3D surface method, value=255), smoothing (uniform, value =4) and fixed thresholding (value=200). The minimum size threshold for organoid detection was 100μm^2^, with no maximum size threshold. Reference levels were calculated to identify organoids of sizes greater than 100um^2^, 1000um^2^, 10,000um^2^ and 15,000um^2^. Images from the 549nm laser were processed for background removal (3D surface method, value=150), smoothing (uniform, value=2) and fixed thresholding (value =30). Proximal CAF detection was performed by using a mask from organoids detected from the 485nm channel with mask type ring with a width of 12um and distance of −8um from the perimeter of the detected organoid. Detection of all CAFs in field was performed by using a mask from organoids detected from the 485nm channel with mask type circle and maximal size (value = 255) to cover the entire area of the field.

After image acquisition, cultures were then subjected to a resazurin reduction potential assay as a final end-point measure for culture viability. Cultures were incubated with 7 µl Resazurin (PrestoBlue Cell viability reagent, ThermoFisher, A13262) for 1 h prior to fluorescence spectrophotometric measurement for the product of resazurin reduction, resorufin by the SpectraMax I3X from one point per well.

### Data Processing PCA

Output data from the assay measures include full well spectrophotometry for turboGFP, mKate2 and resorufin fluorescence measures. Additionally, 71 metrics were acquired from the high content image analysis measures captured from the two imaging platforms (74 metrics total, (Supplementary table 1)). Spectrophotometry measures for turboGFP and mKate2 were each averaged across the 9 points taken per well. All metrics for all co-culture conditions (co-cultures exposed to compound, DMSO vehicle control co-cultures and cycloheximide co-cultures) within each plate were then processed for a non-stringent outlier removal, removing measures outside 3 times the interquartile range from the median measure of each metric for that plate (Performed in R, (R Core Team (2021)). This was followed by a scaling of each metric in a plate-wise manner (Performed in R). All plates were then compared together by PCA using Simca 16.0 software (Umetrics, Sartorius Stedim Biotech), unit variance scaled and performing a 7-fold cross validation. The cross validation was used to calculate a Q^2^ score for each PC to indicate the predictive capacity of the PCA model, and additional PCs were calculated as long as the cumulative Q^2^ score increased, that is, each new PC still contributed to the predictive capacity of the model and unlikely to be modelling arbitrary noise. Compound selection from PCA was performed by calculating the product of PC 1 and 2 from those data that were both positive for those PCs and selecting those compounds with the highest product.

### Data Processing K-means

Data from the screen were subject to a non-stringent outlier removal, removing measures outside 3 times the interquartile range from the median measure of each metric for that plate, then data from co-cultures exposed to test compounds only (not using controls) were plate-wise scaled. These data were modelled together by PCA using Simca 16.0 software (Umetrics, Sartorius Stedim Biotech), unit variance scaled and performing a 7-fold cross validation. The cross validation was used to calculate a Q^2^ score for each PC to indicate the predictive capacity of the PCA model, and additional PCs were calculated as long as the cumulative Q^2^ score increased, that is, each new PC still contributed to the predictive capacity of the model and unlikely to be modelling arbitrary noise. The loadings from all principal components from the PCA that still had explanatory value were acquired. Loadings were unit variance scaled and then used as input data for K-means analysis. The number of clusters to use was calculated by examining the gap statistic algorithm (Performed in R). Clusters were then calculated, and the z-scores (determined from the initial plate-wise scaling) of the metrics measured from co-culture wells belonging to each cluster were examined to interpret phenotype. A cluster with suitable metrics was selected and the top 704 compounds were selected by identifying those data from the selected cluster that were furthest from the origin in an n-dimensional plot of all the PCs that were used in the K-means clustering.

### Traditional selection method for initial screen

Data from the initial screen were processed for a non-stringent outlier removal, removing measures outside three times the interquartile range from the median measure of each metric for that plate (Performed in R). The data for each metric was standardized plate-wise as a fold change of the mean value of the DMSO vehicle control cultures of each plate. Hit compounds were then selected according to the criteria outlined in Table 2.

### Hit Verification Screen

The selected 704 compounds were acquired from CBCS which had been maintained under their storage conditions. Upon receipt, compounds were maintained in DMSO at −20°C. The compounds were ordered for final screening concentrations of 5, 10 and 20 μM. The screening procedure mirrored that of the initial screen in terms of co-culture preparation, plating and plate-map except with the relevant selected compounds.

### OPLS selection from hit verification screen

Data from the initial screen (∼36,000 compounds) and the hit verification screen (704 compounds) were processed for a non-stringent outlier removal, removing measures outside 3 times the interquartile range from the median measure of each metric for that plate (Performed in R). To compare data between the screens, the data for each metric was standardized plate-wise as a fold change of the mean value of the DMSO vehicle control cultures of each plate. A comparison between the fold-change standardization and the plate-wise scaling approach of the initial screen was performed. An OPLS model was trained using data from the initial screen, including all DMSO vehicle control treated co-cultures, all cycloheximide treated co-cultures and the 704 selected compounds selected for the hit verification screen as classifiers. Data were unit-variance scaled and modelled with a seven-fold cross-validation. The data from the hit verification screen was then tested against the model as a prediction set for classification into the three classifiers used in training the model (DMSO, cycloheximide, selected compound). Compounds that were predicted to be either DMSO- or cycloheximide-treated at any concentration were excluded from further selection. Remaining compounds were ranked according to their average predictive score as a test compound across the three tested concentrations.

### Traditional selection from hit verification

Data from the hit verification screen were processed for a non-stringent outlier removal, removing measures outside three times the interquartile range from the median measure of each metric for that plate (Performed in R). The data for each metric was standardized plate-wise as a fold change of the mean value of the DMSO vehicle control cultures of each plate. Hit compounds were then selected according to the criteria outlined in Table 3.

### GNF-5 Dose Response

GNF-5 was acquired from CBCS which had been maintained under their storage conditions. GNF-5 was dissolved in DMSO and stored at −20°C. GNF-5 was used in dose-response assays at concentrations of 20, 10, 5, 2, 1, 0.3, 0.12 and 0.05μM. The screening procedure mirrored that of the initial screen in terms of co-culture preparation, plating and time course. Culture formats included additional biological replicate co-cultures (mT3.tGFP:PSC5mKate2, mT4.tGFP:PSC4.mKate2 and mT5.tGFP:PSC5.mKate2), as well as one tumor organoid mono-culture condition (mT3.tGFP) and three biological replicate mono-cultures of non-tumor murine pancreatic ductal epithelial organoids (mN5, mN11, mN12). The cell dissociation for the non-tumor organoids was identical to that of the tumor organoids as previously described. Since non-tumor organoids cannot be maintained in the reduced media condition and so all these replicates and the mT3.tGFP mono-culture test conditions were maintained in complete organoid media supplemented with 10.5μM Y-27632 and compound or DMSO vehicle control. Data collection mirrored that of previous analyses with the addition of brightfield analyses for mono-cultures analysed by SoftmaxPro 7.0 software to collect percentage covered area data of organoids.

### Therapeutic window calculations

To compare the effect of compounds on tumor cells in co-culture and non-tumor cells in mono-culture, a comparison was performed by comparing the resazurin reduction assay data and the percentage covered area data from the minimax microscope. In the case of tumor co-cultures, the data from the green fluorescence percentage covered area was used, whereas in the non-tumor mono-cultures, the percentage covered area data from the brightfield images was used. To compare these data between culture types (resazurin, green fluorescence, brightfield), the data for each of these metrics was plate-wise rescaled according to the average respective values from the DMSO vehicle controls and the cycloheximide treated controls where a value of 1 was equivalent to the mean DMSO condition and 0 equivalent to the mean cycloheximide condition. Additionally, a similar approach was taken with the PSC percentage covered area in co-cultures. These data were then modelled for dose response in GraphPad Prism (version 9, GraphPad Software) using the non-linear regression variable slope analysis (four parameters) function.

## Supporting information

Description of CBCS Screen Library

## Acknowledgements

We would like to thank Professor David Tuveson, Cold Spring Harbor Laboratory (New York, USA) for donation of the cell lines used in experiments. We acknowledge the common flow cytometry platform at Umeå university (Flow@CliMi) for providing access to and assistance with flow cytometry experiments. We are grateful also for the help and support from CBCS.

## Supplementary Figure legends

**Supplementary Figure 1**

**FACS analysis.** Gating strategy for flow cytometry for cancer activated fibroblast (CAF) subtype analysis and fluorescence-activated cell sorting (FACS) of CAFs from tumor-CAF co-cultures. Cells are gated for size to avoid cellular debris (Forward scatter (FSC) vs. side scatter (SSC)), singlets by DAPI (width vs. area) and for live cells by live/dead stain (APC-Cy7 area vs. SSC). For CAF subtypes, cultures are gated for PDGFR-b for CAF identity and analysed for alpha smooth muscle actin expression (α-SMA, myofibroblastic CAF marker) and for interleukin-six (Il6, inflammatory CAF marker). For FACS sorting, CAFs are gated by PDGFR-b for sorting. For each marker, gates were determined by examination of cells negative for marker expression, such as mono-cultured tumor cells for PDGFR-b, α-SMA and Il-6.

**Supplementary figure 2**

**Plasmid construct maps.** Plasmid maps for constitutive fluorescent protein expression in cells by lentiviral transfection for **(A)** turbo-GFP and **(B)** mKate2 expression; both driven by translation elongation factor 1 alpha (EF1A) and with neomycin (Neo) resistance.

**Supplementary figure 3**

**Drug screen using Arrayscan imaging platform. (A)** Visual representation of analysis from Arrayscan imaging. The top row displays pseudo-colored images from the Arrayscan microscope as individual channels and a merge, and underneath displays the output of object detection for all organoids (outlined in green), proximal CAFS (red, detection radius outlined in red line; origin is detected organoids) and distal CAFs (yellow, detection radius outlined in yellow lines; origin is detected organoid). Additionally, Organoids were analysed for size; highlighted here are “extra-large” organoids (>15,000μm^2^, filled green). **(B)** Plate-map used for the screening process in 384 well-plates. Per plate; 352 wells were used for testing compounds in co-cultures, and the remaining 16 used for control wells. Control wells included 8 co-cultures, 8 tumor organoid mono-cultures, 4 mono-cultures of PSCs, and 4 matrigel-only background controls each treated with reduced media containing DMSO vehicle control. Additionally, 4 mono-cultures of tumor organoids were maintained in complete organoid feeding medium and 4 co-cultures were maintained in reduced medium containing cycloheximide (table 1).

**Supplementary Figure 4**

**Principal component analysis (PCA) features.** Principal component loadings of the 74 metrics used in building the PCA for the initial screen for **(A)** principal component (PC) 1 and **(B)** PC 2. Loadings are arranged as a waterfall plot in ascending order. **(C, D)** An analysis of the 74 metric contributions to different areas of the PCA by quadrant. Here, the average of the 74 metrics of highlighted data (red) are compared to the average of metrics across the screen and presented as waterfall plots in ascending order for compounds found in **(C)** the bottom left and **(D)** top right of the PCA plot. Data negative in the comparison (orange bars) indicate that highlighted cultures (red) had lower measures on average for that metric, whereas data positive in the comparison (orange bars) indicates that the highlighted cultures (red) had larger measures for that metric on average.

**Supplementary figure 5**

**Fit scores for principal components (PC) in principal component analysis (PCA) and k-means clustering models.** Cumulative goodness of fit (R^2^, R2X(cum), green) explains the proportion of variance in the dataset incorporated into the model and the cumulative predictive capacity of the model (Q^2^, Q2(cum), blue) is determined by 7-fold cross-validation of the models. Additional PCs are calculated so long as Q^2^ increases to avoid modelling noise. **(A)** The PCA model used for PCA selection procedure was built incorporating data from control wells and contains 11 PCs. **(B)** The PCA model used for dimensionality reduction for later K-means clustering did not include data from these controls and contains 13 PCs. **(C)** Gap-statistic for optimal number of k-means clusters; a gap-statistic is calculated for each number of clusters in a k-means model. A local maxima at 14 clusters is identified. **(D)** Clusters determined by K-means clustering algorithm (14 clusters) are plotted in a PCA plot and pseudo-colored according to cluster identity. **(E)** Waterfall boxplot (order determined by median) of the z-score of the 74 metrics measured from each culture in the initial screen for cluster 14.

**Supplementary figure 6**

**Correlation between dataset standardizing approaches in PCA and OPLS models. (A)** Correlation of loadings for principal component (PC) 1 (left) and PC2 (right) between principal component analysis models using data from the initial screen that has been standardized by two different methods. The x-axis represents the loadings from data standardized by plat-wise scaling, whereas the y-axis represents the loadings from the same data standardized in terms of fold change of vehicle control co-cultures. Each point represents one of the 74 metrics measured and modelled in the two PCAs. **(B)** Summary of fit for OPLS model built for the hit verification analyses. Cumulative goodness of fit (R^2^, R2X(cum), green) explains the proportion of variance in the dataset incorporated into the model and the cumulative predictive capacity of the model (Q^2^, Q2(cum), blue) is determined by 7-fold cross-validation of the models. The model contains two components. **(C)** Receiver-operating characteristic of the OPLS model examining the training data-set used to build the model.

**Supplementary figure 7**

**Functional protein association network**. **(A)** RNA expression levels indicate low cancer specificity being ABL1 overexpressed in multiple cancer types such as lung, colorectal, leukemia and pancreatic cancer. **(B)** ABL1 expression is enriched in endocrine, ductal, endothelial, mesenchymal, and immune cells (https://www.proteinatlas.org/ENSG00000097007-ABL1/tissue+cell+type/pancreas). **(C)** Protein-protein interaction networks functional enrichment analysis using String database (https://string-db.org) reveal ABL1 clusters together with ACTA2 (α-SMA), ACTA1, CTNNB1, BCL9 and APC amongst other interactors.

**Supplementary figure 8**

**Object detection correlation from different channels of a high content imaging platform.** Correlation of object detection for organoid covered area of mono-cultures of fluorescent tumor organoids between images gathered by brightfield (y-axis) and fluorescence images (x-axis). Each point represents the results of each analysis for the same culture. Cultures examined are mono-cultures of tumor organoids exposed to DMSO vehicle control (n=8), GNF-5 (n=8, one per eight concentrations 0.05-20μM) and cycloheximide (n=4, 250ng/ml). Plotted is the line of best fit (black line, linear regression) and 95% confidence intervals (dotted lines).

**Supplementary figure 9**

**ABL1 expression and localization in human tissue and cells.** Expression clustering and correlation between ABL1 and relevant transforming growth factors involved in myofibroblast differentiation and ECM organization was found in cluster 57 of pancreatic fibroblasts at the Human Protein Atlas (https://www.proteinatlas.org/ENSG00000097007-ABL1).

