## Supplementary Table 1 for "High throughput screen in a co-culture model to uncover therapeutic strategies to potentiate the cancer-inhibiting properties of the tumor-stroma in pancreatic cancer"

### Supplementary Table 1 – Metrics modelled in analyses

| METRIC Nº (#) | NAME | TYPE OF MEASURE* | RAW OR CALCULATED | CALCULATION |
|---|---|---|---|---|
| **1** | tGFP Fluorescence | S | Raw | |
| **2** | mKate2 Fluorescence | S | Raw | |
| **3** | Resazurin | S | Raw | |
| **4** | Organoid count | M | Raw | |
| **5** | Organoid Covered Area (%) | M | Raw | |
| **6** | Organoid average area (um2) | M | Raw | |
| **7** | Organoid average roundness | M | Raw | |
| **8** | Organoid average intensity | M | Raw | |
| **9** | Organoid average total intensity | M | Raw | |
| **10** | Organoid total intensity | M | Raw | |
| **11** | CAF count | M | Raw | |
| **12** | CAF covered area (%) | M | Raw | |
| **13** | CAF Average Area (um2) | M | Raw | |
| **14** | CAF average roundness | M | Raw | |
| **15** | CAF average Intensity | M | Raw | |
| **16** | CAF average total intensity | M | Raw | |
| **17** | CAF Total intensity | M | Raw | |
| **18** | Proximal CAF Count | M | Raw | |
| **19** | proximal CAF average proportion of CAFs | M | Raw | |
| **20** | proximal CAF covered area (%) | M | Raw | |
| **21** | proximal CAF average Area (um2) | M | Raw | |
| **22** | proximal CAF average roundness | M | Raw | |
| **23** | proximal CAF average Intensity | M | Raw | |
| **24** | proximal CAF average total intensity | M | Raw | |
| **25** | proximal CAF Total Intensity | M | Raw | |
| **26** | Distal CAF Count | M | Raw | |
| **27** | Distal CAF proportion of CAFs | M | Raw | |
| **28** | Distal CAF average area (%) | M | Raw | |
| **29** | Distal CAF average area (um2) | M | Raw | |
| **30** | Distal CAF average roundness | M | Raw | |
| **31** | Distal CAF average Intensity | M | Raw | |
| **32** | Distal CAF average total intensity | M | Raw | |
| **33** | Distal CAF Total Intensity | M | Raw | |
| **34** | Organoid Count | A | Raw | |
| **35** | Organoid count (Small, <1,000um2) | A | calculated | Organoid count*Organoid proportion (#34*#39) |
| **36** | Organoid count (medium, 1,000<x<10,000um2) | A | calculated | Organoid Count > 10,000 - Organoid Count xl (#unlisted-#38) |
| **37** | Organoid count (large, 10,000<x<15,000 um2) | A | Calculated | Organoid count - Organoid count (s, m, xl) (#34-#35-#36-#38) |
| **38** | Organoid count (extra-large, >15,000um2) | A | Raw | |
| **39** | Organoid proportion (small) | A | Raw | |
| **40** | Organoid proportion (medium) | A | Calculated | Organoid count (m)/Organoid count (#36/#34) |
| **41** | Organoid proportion (large) | A | Calculated | Organoid count (l)/ Organoid count (#37/#34) |
| **42** | Organoid proportion (extra-large) | A | Calculated | Organoid count xl / total organoid count (#38/#34) |
| **43** | Organoid total area | A | Raw | |

| | | | | |
|---|---|---|---|---|
| **44** | Organoid area (extra large, >15,000um2) | A | Raw | |
| **45** | Organoid area (small-Large, <15,000um2) | A | Calculated | Organoid total area - Organoid area xl, (#43-#44) |
| **46** | Proportion area (small-large, <15,000 um2) | A | Calculated | Organoid area (s-l)/Organoid total area (#45/#43) |
| **47** | Proportion area (extra-large, >15,000um2) | A | Calculated | Organoid area (xl)/Organoid total area (#44/#43) |
| **48** | CAF total count | A | Raw | |
| **49** | Proximal CAF count | A | Raw | |
| **50** | Distal CAF count | A | Calculated | CAF total count - Proximal CAF count (#48-#49) |
| **51** | Proximal CAF proportion | A | Calculated | Proximal CAF count/CAF total count (#49/#48) |
| **52** | Distal CAF proportion | A | Calculated | Distal CAF count/CAF total count (#50/#48) |
| **53** | CAF per Organoid | A | Calculated | CAF total count/ Organoid count (#48/#34) |
| **54** | Proximal CAF per Organoid | A | Calculated | Proximal CAF count/Organoid count (#49/#34) |
| **55** | Distal CAF per Organoid | A | Calculated | Distal CAF count/Organoid count (#50/#34) |
| **56** | Proximal CAF per area Organoid | A | Calculated | Proximal CAF count/ Organoid total area (#48/#43) |
| **57** | CAF total area | A | Raw | |
| **58** | Proximal CAF area | A | Raw | |
| **59** | Distal CAF area | A | Calculated | CAF total area - proximal CAF area (#57-#58) |
| **60** | Proximal CAF area proportion | A | Calculated | Proximal CAF area/CAF total area (#58/#57) |
| **61** | Distal CAF area proportion | A | Calculated | Distal CAF area/ CAF total area (#59/#57) |
| **62** | Proximal CAF area per Organoid area | A | Calculated | Proximal CAF area/Organoid total area (#58/#43) |
| **63** | Distal CAF area per non-Organoid area | A | Calculated | Distal CAF area/ (total well area-Organoid total area) (#59/(#unlisted-#43) |
| **64** | Organoid total area (um2) | M | Calculated | Organoid count * Organoid average area (#4*#6) |
| **65** | CAFs per Organoid | M | Calculated | CAF count/Organoid count (#11/#4) |
| **66** | CAF total area (um2) | M | Calculated | CAF count * CAF average area (#11*#13) |
| **67** | CAF area per Organoid area | M | Calculated | CAF total area/Organoid total area (#66/#64) |
| **68** | Proximal CAF count/Organoid count | M | Calculated | Proximal CAF count / Organoid count (#18/#4) |
| **69** | Proximal CAF total area (um2) | M | Calculated | Proximal CAF count * Proximal CAF average area (#18*#21) |
| **70** | Proximal CAF area per Organoid area | M | Calculated | Proximal CAF total area/Organoid total area (#69/#64) |
| **71** | Distal CAF count per Organoid | M | Calculated | Distal CAF count/Organoid count (#26/#4) |
| **72** | Distal CAF total area (um2) | M | Calculated | Distal CAF count * Distal CAF average area (#26*#29) |
| **73** | Distal CAF density | M | Calculated | Distal CAF area/ field area-Organoid area) (#72/(#unlisted-#64)) |
| **74** | Proximal CAF density per Distal CAF density | M | Calculated | (Proximal CAF area/Organoid total area)/Distal CAF density (#70/#73) |

* Type of measure: S, spectrophotometry; M, Minimax microscopy; A, Arrayscan Microscopy
