## Supplementary material for "High throughput screen in a co-culture model to uncover therapeutic strategies to potentiate the cancer-inhibiting properties of the tumor-stroma in pancreatic cancer": Description of CBCS Screen Library

**Primary set screen:**

The libraries of compounds applied in this screening campaign consists of the primary screening set at CBCS of 33084 compounds and the CBCS annotated set of 5488 compounds. The compounds were donated by various biotech companies and originates from both in-house and commercial sources. Compounds included in the primary screening set were selected to represent a diverse selection of a larger set of >350,000 compounds, while keeping a certain depth to allow crude structure–activity relationship studies. The selection was also biased towards lead-like and drug-like profiles with regards to molecular weight, hydrogen bond donors/acceptors and log P. The CBCS annotated library includes known kinase- protease- phosphatase- and GPCR-inhibitors, epigenetic- autophagy- and Wnt-modulators, nuclear receptor and ion channel-ligands, redox compounds and a set of FDA & EMA approved drugs. Compound stock solutions at 10 mM in DMSO are stored frozen at approximately −20 °C in individual capped tubes in REMP 96 Storage Tube Racks. The racks are stored in a REMP Small-Size Store, which allows cherry picking while the solutions are still frozen to minimize repetitive freeze-thaw cycles. For screening purposes, the compound solutions have been replicated from the REMP racks to Labcyte 384 LDV plates (LP-0200) and then further into Labcyte 1536 HighBase plates (LP-03730) to enable dispensing using acoustic liquid.

Facts:

Size of Primary screening set:

| | |
|---|---|
| 2010 | 5K |
| 2011-2013 | 10K |
| 2014-2015 | 17K |
| 2016-2020 | 30K |
| 2021- | 37K |

**Storage**

The screening compounds (10mM in DMSO) are stored at -18°C deg

Stock solutions and plates (384) are stored in micro vials at -18°C deg

Solid compounds are stored in glass vials at room temperature.

**Retest**

Confirmation of activity is made from the stock solutions using an acoustic liquid handler. The dispensing is done by spotting volumes (2-300nL) to a test plate for further dilution

Annas comments (Libraries available 2019)

| **Libraries available** | **Antal cpds** | **Profile** |
|---|---|---|

| | | |
|---|---|---|
| **Primary Screening set (33K)** | 33084 | |
| *33K subset LCBKI Diverse (div1, frag, LL, DL, addon) | 6138 | Diverse |
| *33K subset SFG (GPCR) | 1986 | Compounds similar to GPCR inhibitors |
| *33K subset SFK (kinases) | 1838 | Compounds similar to kinase inhibitors |
| *33K subset Analyticon | 998 | Natural product inspired |
| *33K subset arach pathway | 1280 | Targets the arachidonic pathway |
| *33K subset ChemBridge_LCBU part 1 | 9806 | Diverse |
| *33K subset Nucleoside Library | 192 | Nucleosides |
| *33K subset KDex_PPI | 1008 | Protein-protein interaction |
| *33K subset KDex_Synergy | 1987 | Diverse |
| *33K subset KDex_Elite | 2306 | Diverse |
| *33K subset KDex_Acids | 816 | Carboxylic acids |
| *33K subset KDex_kinase_targeted | 1313 | Compounds similar to kinase inhibitors |
| *33K subset KDex_Zn_binders | 212 | Predicted Zn-chelators |
| *33K subset Asinex Macrocycles | 220 | Macrocyclic compounds |
| *33K subset Blue dept | 2951 | Diverse, sp3 enriched |
| *33K subset DOLE_Minguez | 33 | Novel spirocyclic scaffolds in the interface between "druglike" molecules and natural products |
| **Annotated Libraries** | 5488 | |
| Prestwick chemical lib | 1200 | FDA, EMA approved drugs |
| Tocris_mini known tool cpds | 1120 | Tool compounds |
| NIH clinical collection | 95 | Otava_nucleobase_derivatives |
| SelleckChem known tool cpds | 990 | Tool compounds |
| ENZO_known_redox | 84 | Redox compounds |
| ENZO_known_epigenetics | 43 | Epigenetic modulators |
| ENZO_known_autophagy | 94 | Autophagy modulators |
| ENZO_known_nuclear_receptors | 76 | Nuclear receptor ligands |
| ENZO_known_ionchannels | 71 | Ion channel ligands |
| ENZO_known_protease inh | 53 | protease inhibitors |
| ENZO_known_phosphatase inh | 33 | phosphatase inhibitors |
| ENZO_known_WNT | 75 | WNT modulators |
| KDex_known_kinase_inhibitors | 193 | Kinase inhibitors |
| BioMol_orphan ligand lib | 84 | Description on next sheet |
| BioMol_Endocannabinoid lib | 41 | Description on next sheet |
| BioMol_Neurotransmitter lib | 714 | Description on next sheet |
| BioMol_Nuclear receptor ligand lib | 76 | Description on next sheet |
| Selleck_known_kinase_inhibitors | 378 | Kinase inhibitors |
| SGC_bromodomain_probes | 45 | |
| AZ_pharmacology_toolbox | 23 | |

List of annotated libraries ordered

25__Prestwick chemical lib
26__Tocris_mini known tool cpds
27__NIH clinical collection_otava

28__SelleckChem known tool cpds
29__ENZO_known_redox
30__ENZO_known_epigenetics
31__ENZO_known_autophagy
32__ENZO_known_nuclear_receptors
33__ENZO_known_ionchannels
34__ENZO_known_protease inh
35__ENZO_known_phosphatase inh
36__ENZO_known_WNT
38__BioMol_orphan ligand lib
39__BioMol_Endocannabinoid lib
40__BioMol_Neurotransmitter lib
41__BioMol_Neurotransmitter lib*Adrenergics
42__BioMol_Neurotransmitter lib*Dopaminergics
43__BioMol_Neurotransmitter lib*Serotonergics
44__BioMol_Neurotransmitter lib*Opioids (sigma ligands)
45__BioMol_Neurotransmitter lib*Cholinergics
46__BioMol_Neurotransmitter lib*Histaminergics (Melatonin ligands)
47__BioMol_Neurotransmitter lib*Ionotropic Glutamatergics
48__BioMol_Neurotransmitter lib*Metabotropic Glutamatergics
49__BioMol_Neurotransmitter lib*GABAergics
50__BioMol_Neurotransmitter lib*Purinergics (Adenosines)
51__BioMol_Nuclear receptor ligand lib
52__Selleck_known_kinase_inhibitors
53__SGC_bromodomain_probes
54__AZ_pharmacology_toolboxAdditional libraries:

Descriptions of the content and propose of the libraries can be found at Scilifelab Compound Center (https://compoundcenter.scilifelab.se/).

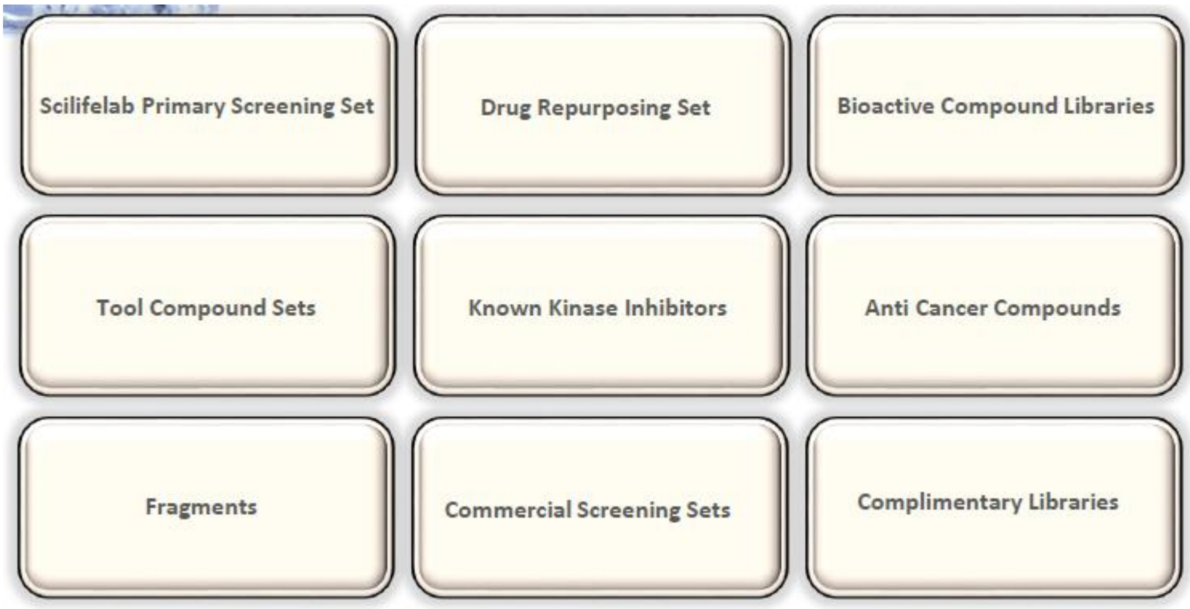


Compound center descriptions:

<u>Prestwick</u>

1200 FDA & EMA approved drugs (current or withdrawn). A diversified marketed drugs library designed for repurposing/repositioning with known bioavailability and safety in humans.

Tocris mini

A library of 1120 biologically active compounds from the Tocris catalog. Covers a wide range of pharmacological targets and research areas.

Enzo known bioactives

529 compounds. Includes compounds that affect most cellular processes and drug target classes. Libraries available: Epigenetics (43), Redox (84), Phosphatase inhibitors (33), Autophagy inducers (71), Autophagy inhibitors (23), Protease inhibitors (53), Nuclear receptor ligand (76)

Sellec known bioactives

A unique collection of 990 bioactive compounds for screening and high content screening

SGC Bromodomain probes

A library of 40 epigenetic chemical probes.

AZ Pharmacological Tool Box

23 compounds with optimized pharmacological properties made available for preclinical research to explore novel disease biology and advance scientific knowledge.

SelleckChem Known Kinase Inhibitors Set

378 known kinase inhibitors available

BioMol (Enzo)Neurotrasmittor Set

700 CNS receptor ligands. Ideal for screening or identifying recombinant orphan G protein-coupled receptors, target validation, secondary screening, validating new assays, and for routine pharmacological applications. Includes 13 classes of receptor ligands.
