## Supplementary figures and images for "High throughput screen in a co-culture model to uncover therapeutic strategies to potentiate the cancer-inhibiting properties of the tumor-stroma in pancreatic cancer"

### Supplementary figure 1

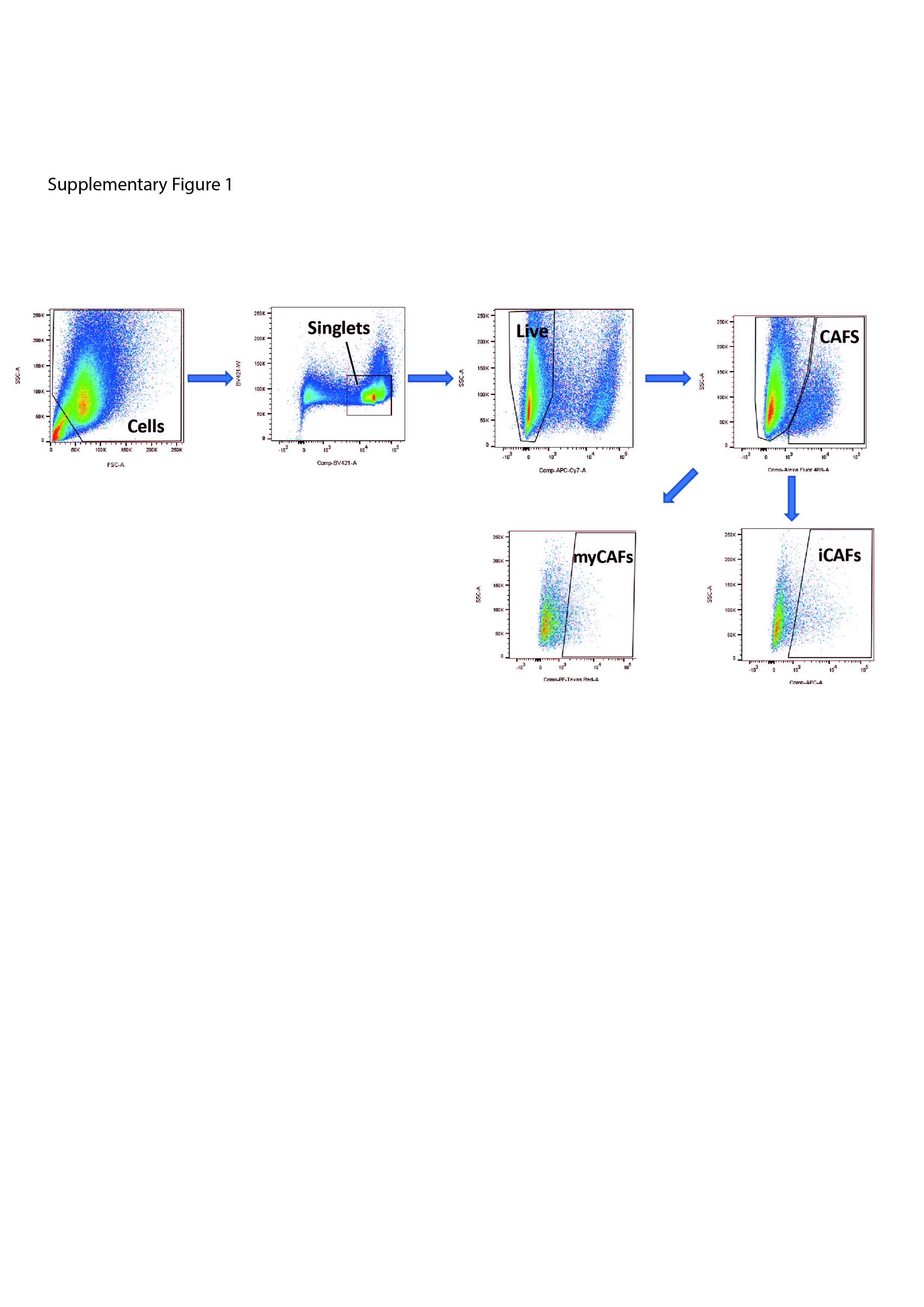

### Supplementary figure 2

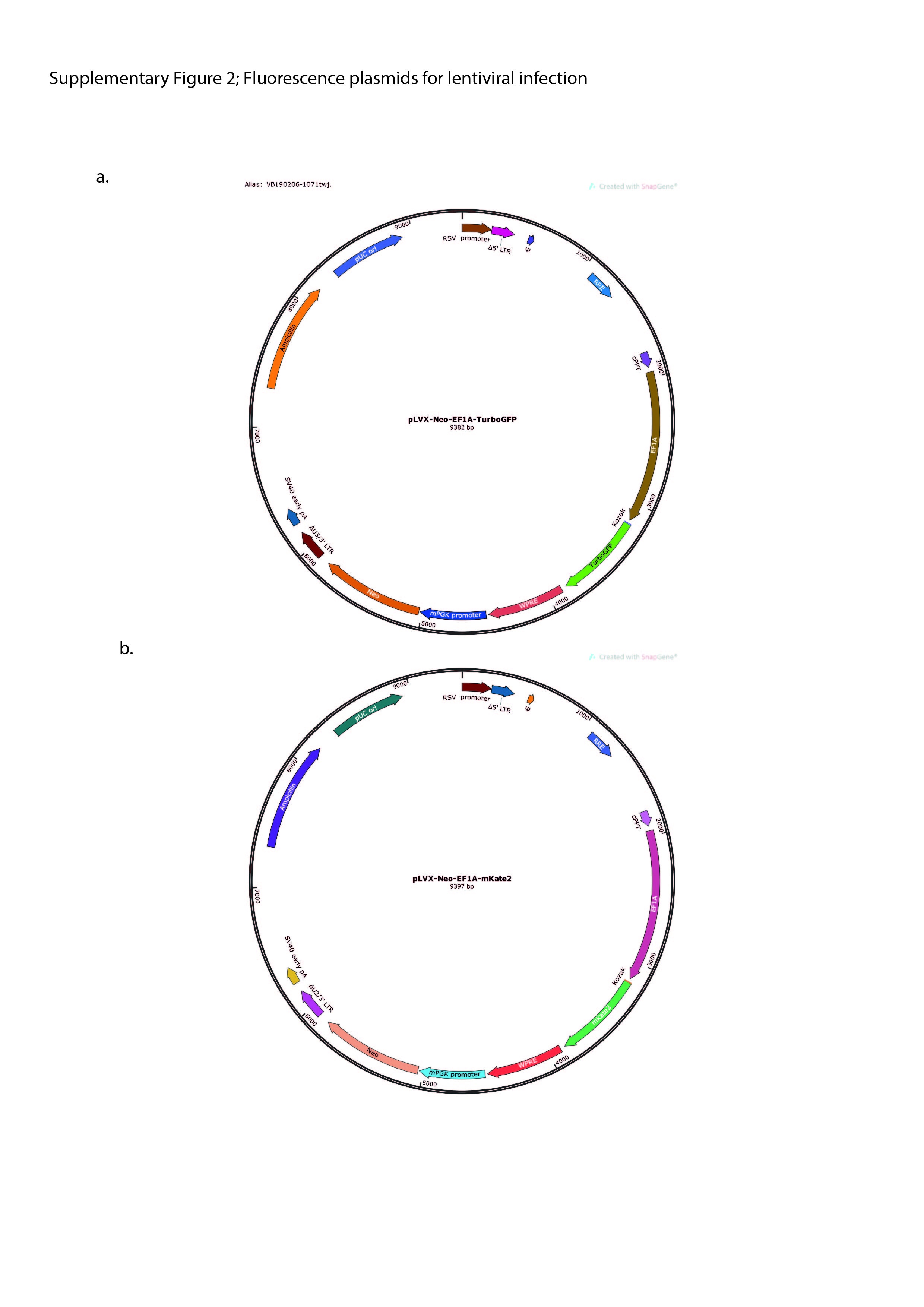

### Supplementary figure 3

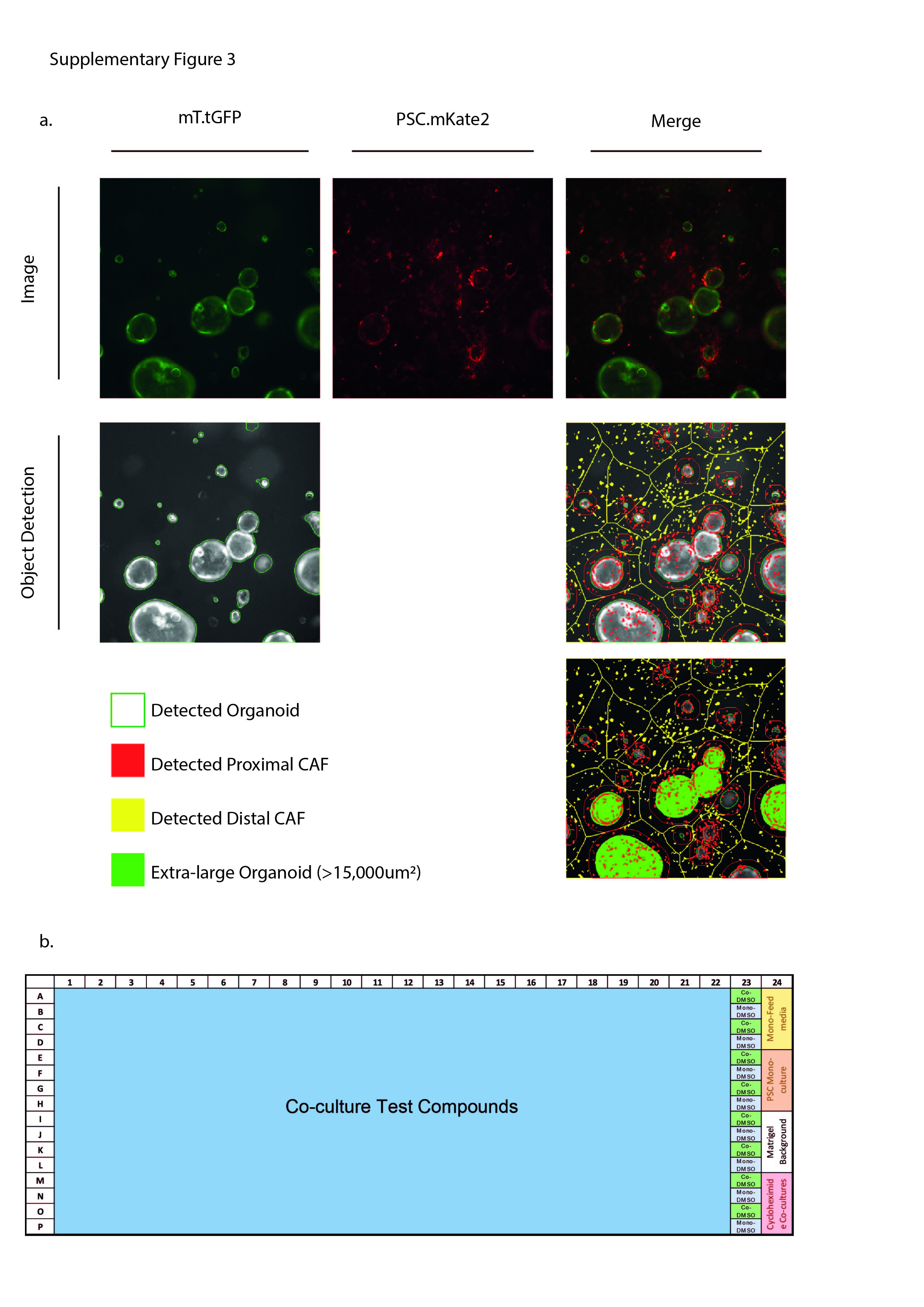

### Supplementary figure 4

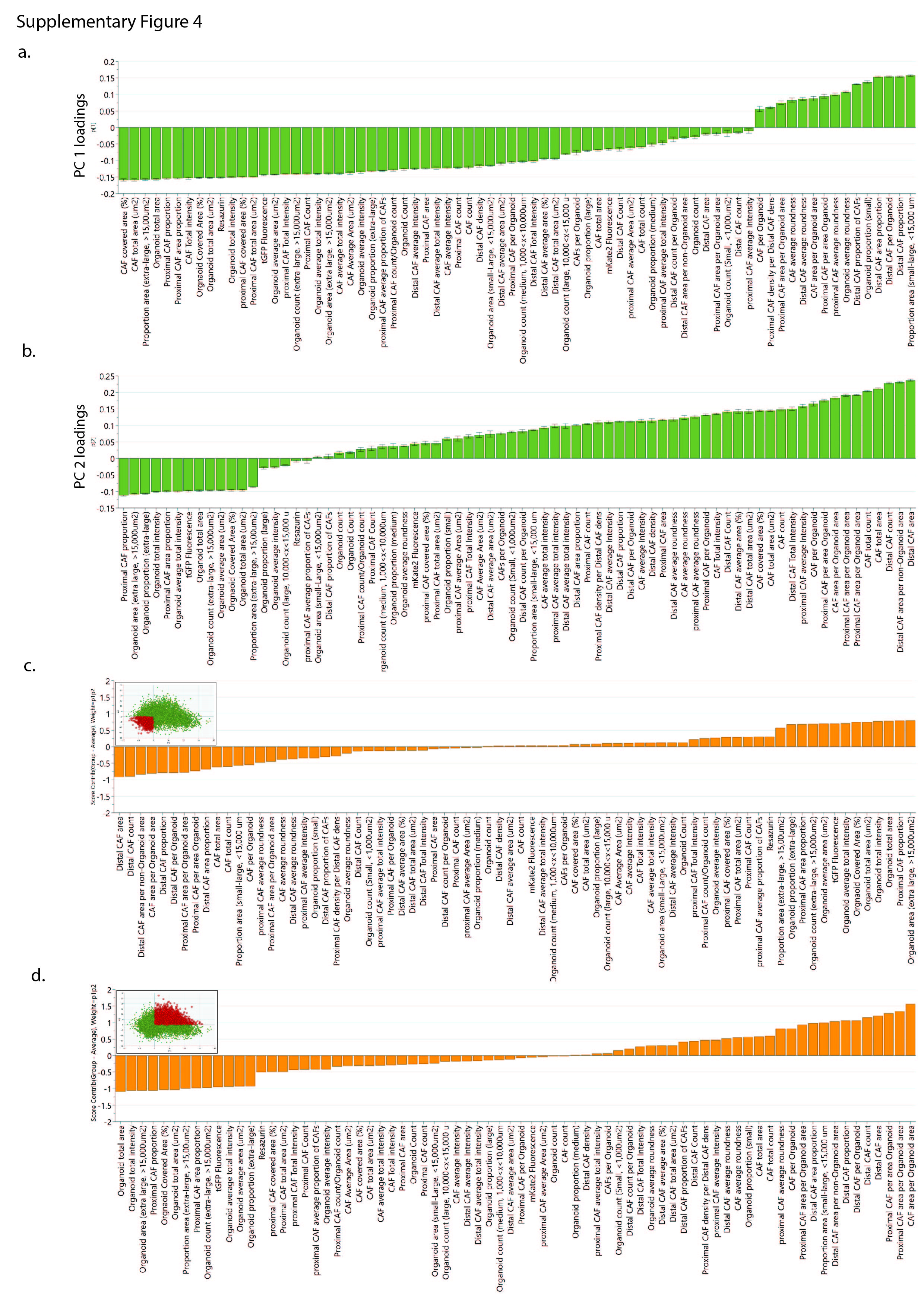

### Supplementary figure 5

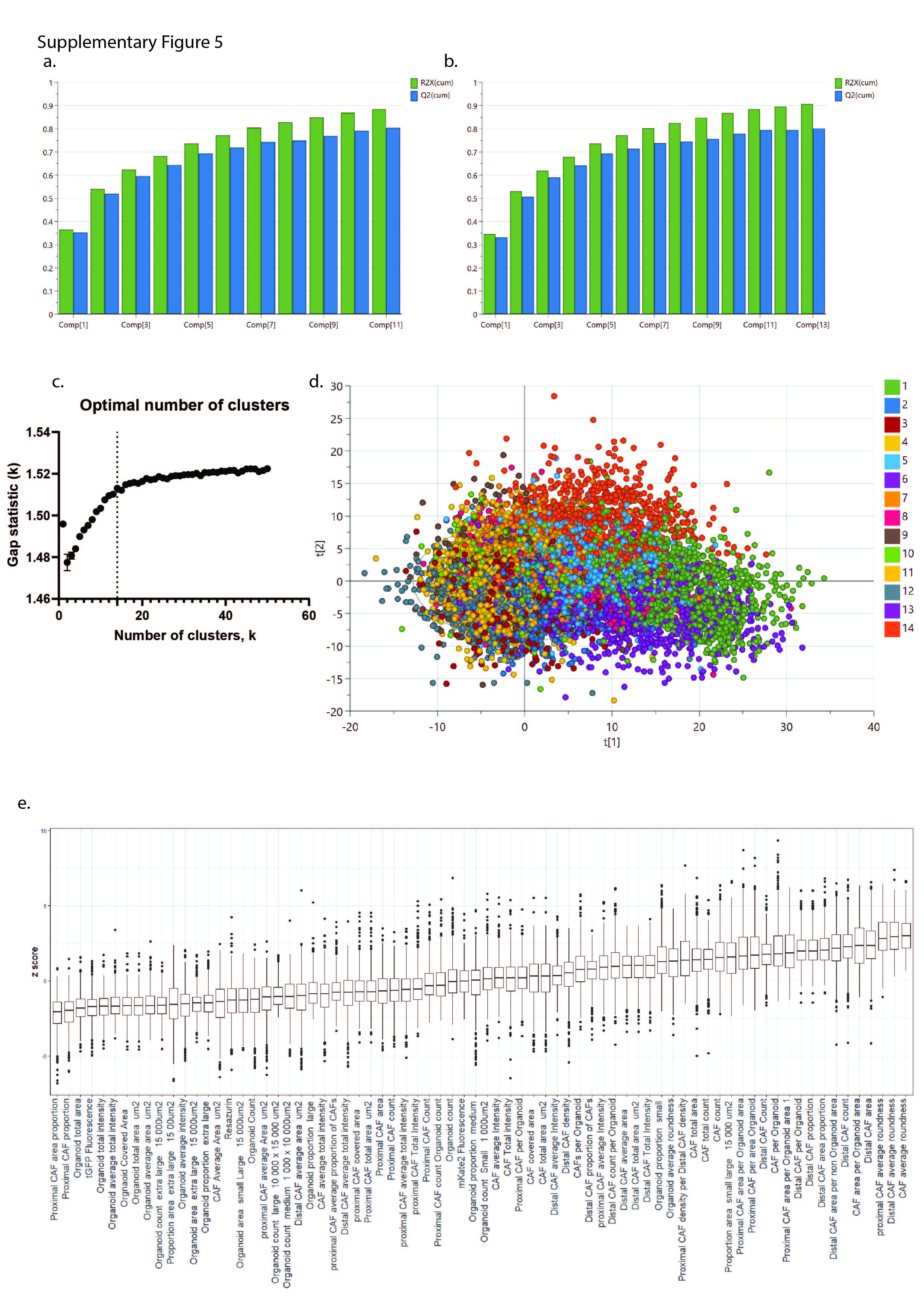

### Supplementary figure 6

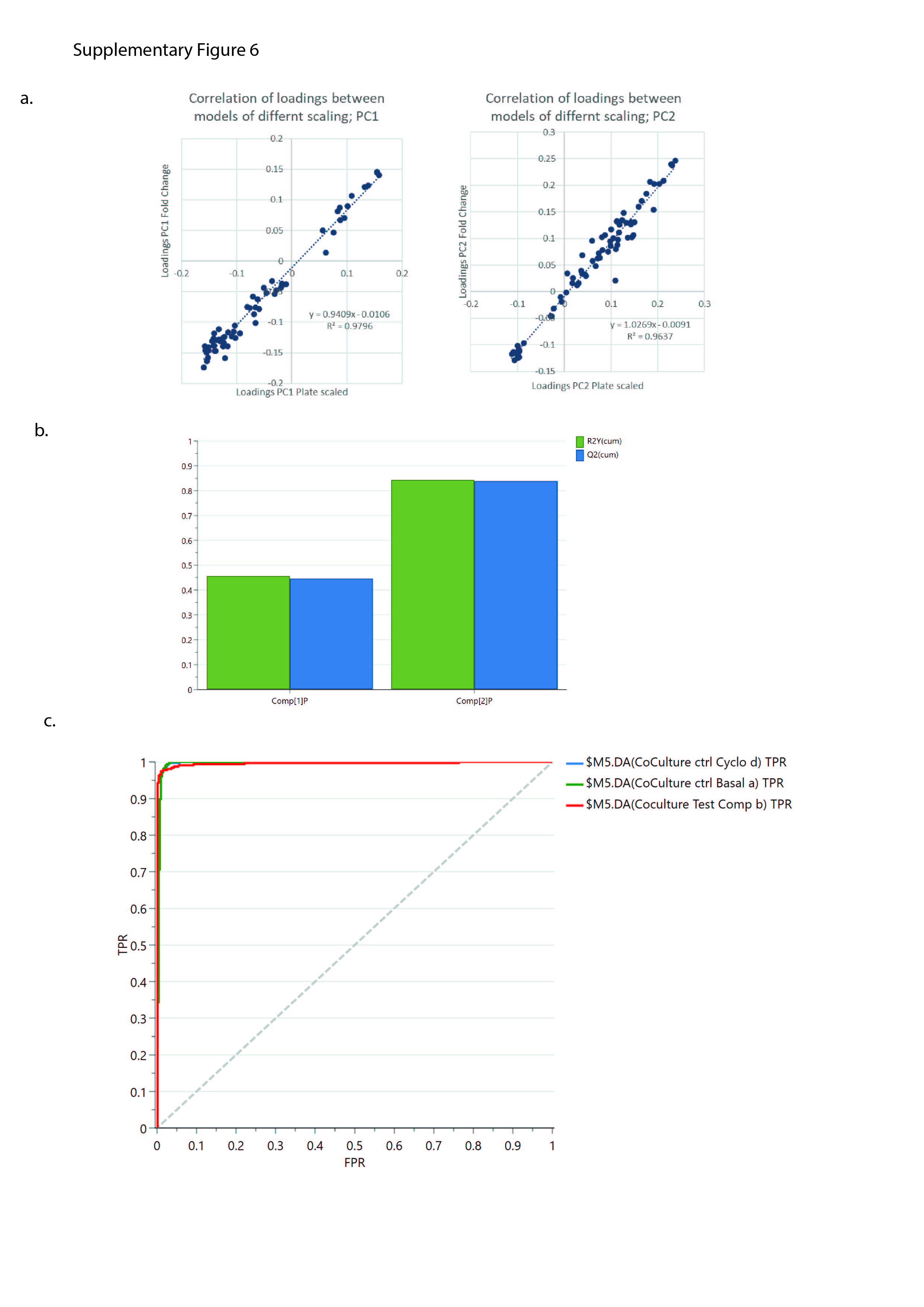

### Supplementary figure 7

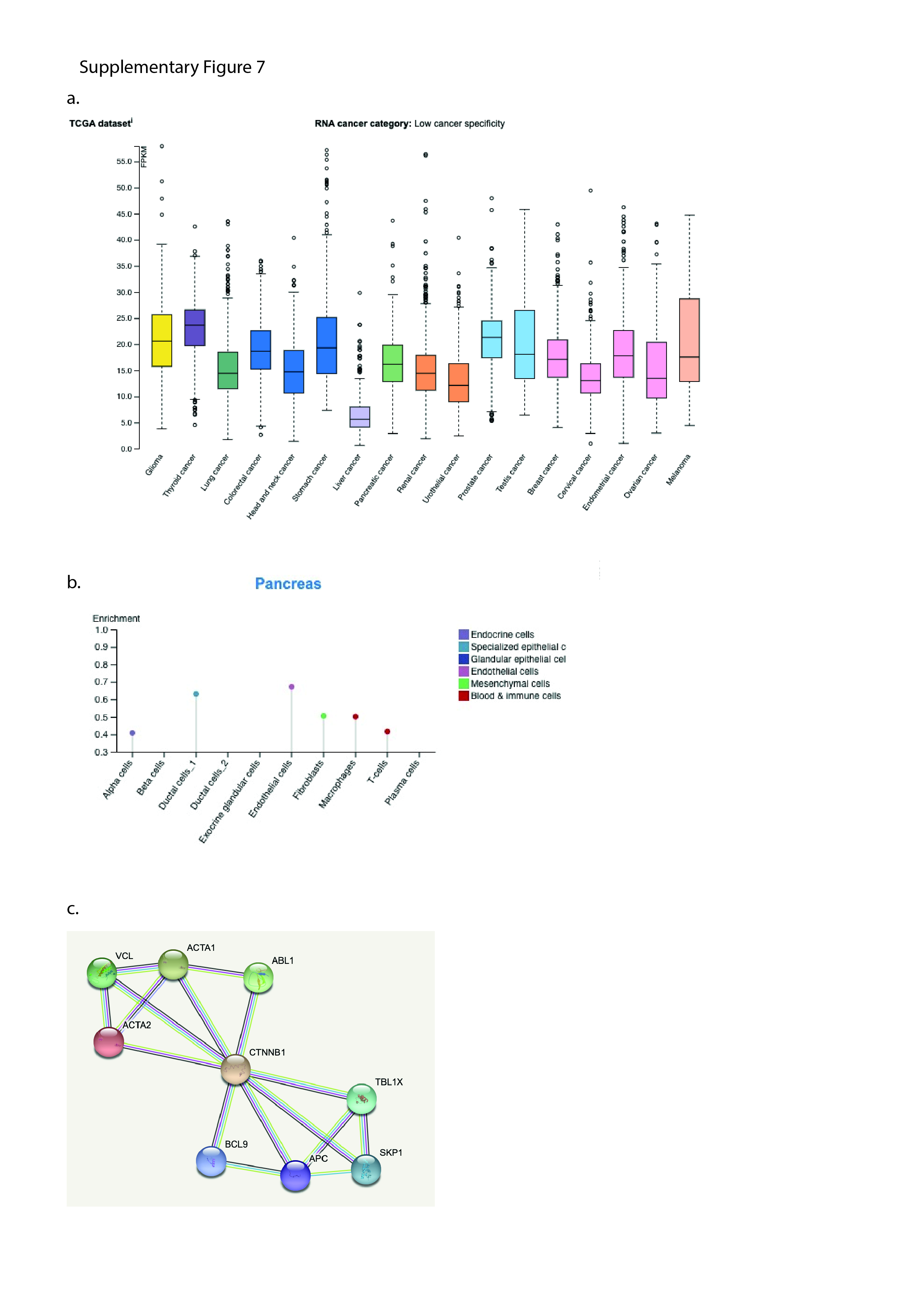

### Supplementary figure 8

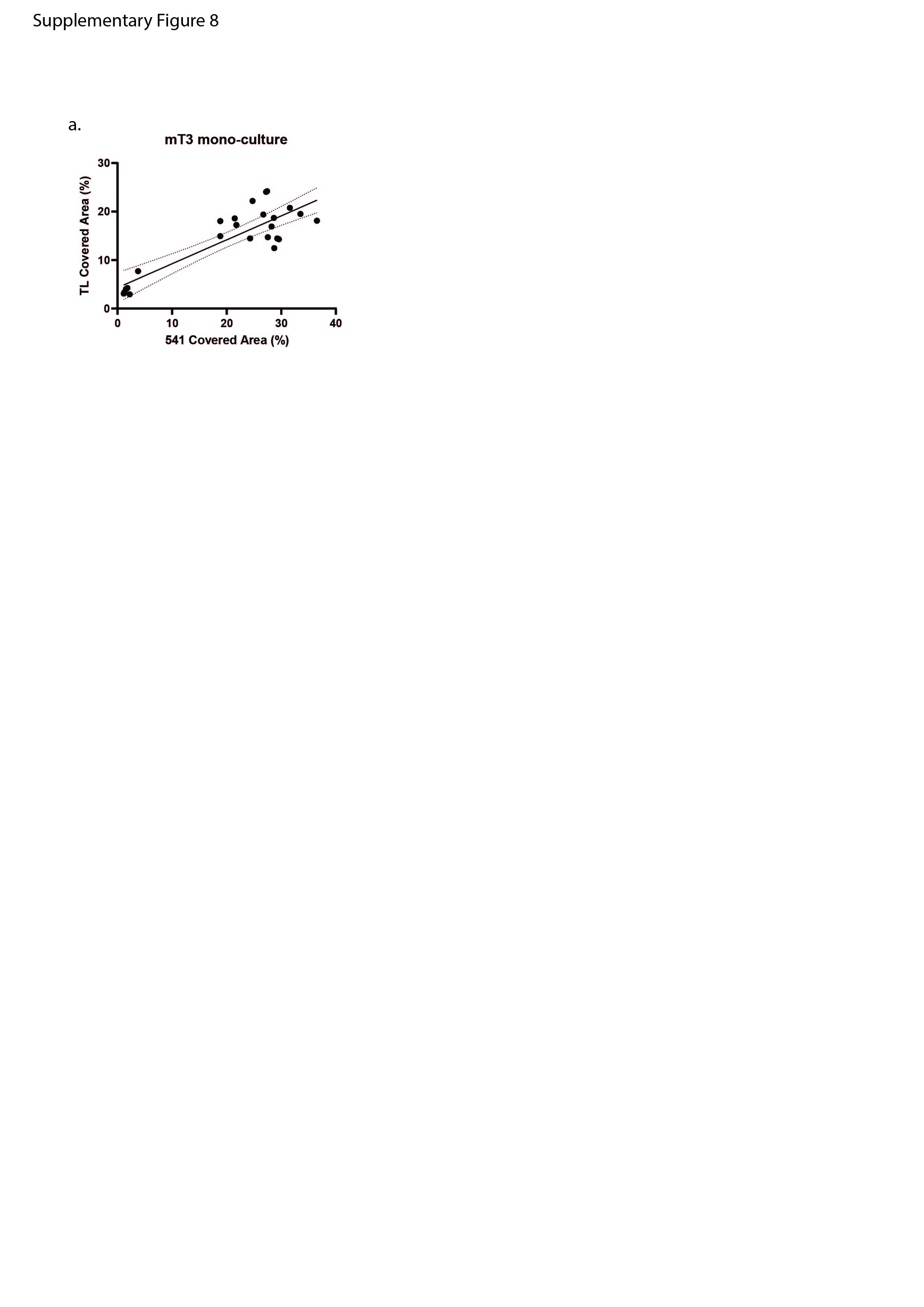

### Supplementary figure 9

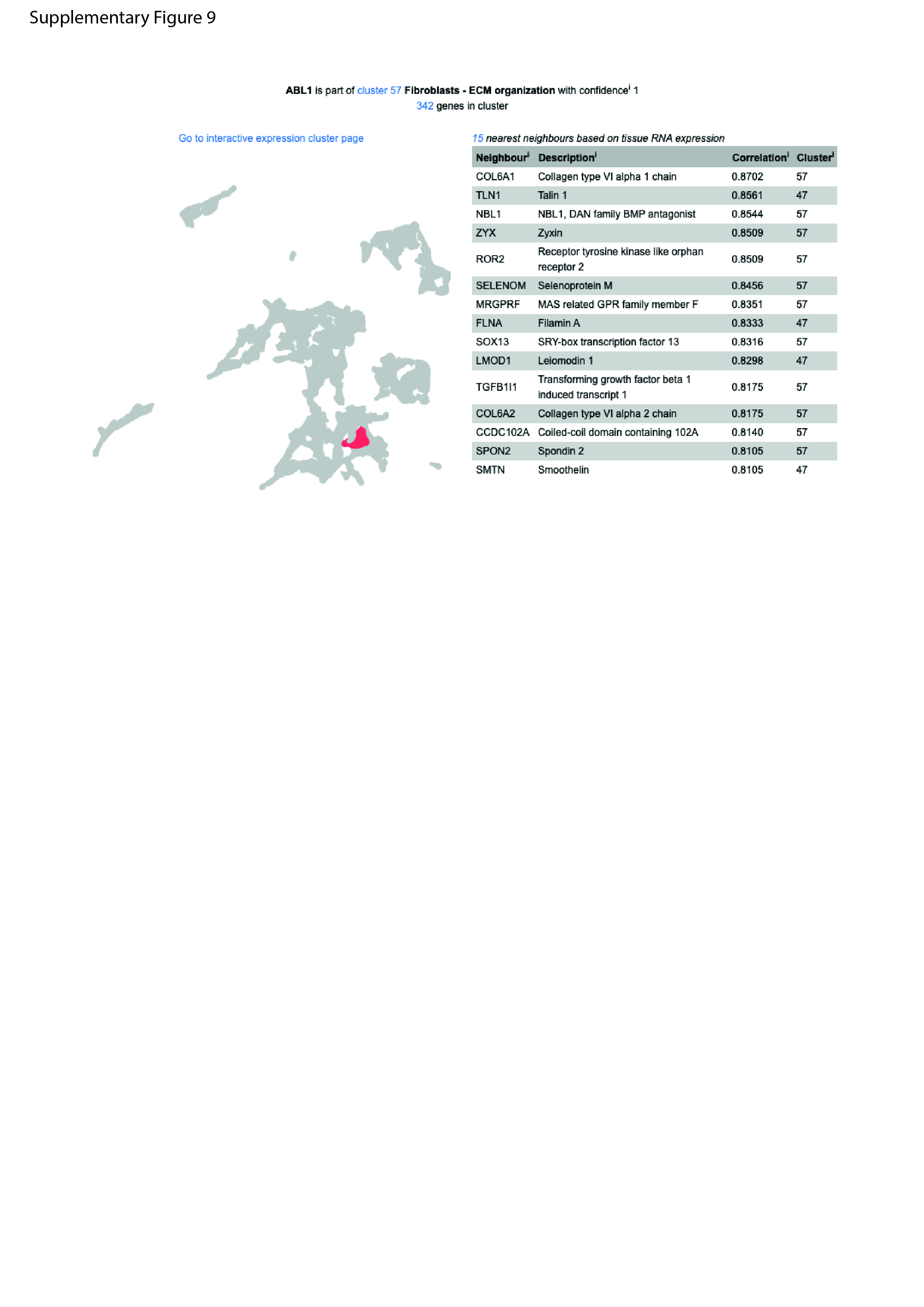
